# Antiviral capacity of the early CD8 T-cell response is predictive of natural control of SIV infection

**DOI:** 10.1101/2023.10.13.562306

**Authors:** Bharadwaj Vemparala, Vincent Madelain, Caroline Passaes, Antoine Millet, Véronique Avettand-Fenoel, Ramsès Djidjou-Demasse, Nathalie Dereuddre-Bosquet, Roger Le Grand, Christine Rouzioux, Bruno Vaslin, Asier Sáez-Cirión, Jérémie Guedj, Narendra M. Dixit

**Affiliations:** Department of Chemical Engineering, Indian Institute of Science, Bengaluru, India; Université Paris Cité, IAME, INSERM, F-75018 Paris, France; Institut Pasteur, Université Paris Cité, Viral Reservoirs and Immune Control Unit, Paris, France; CEA, Université Paris-Saclay, INSERM U1184, Immunology of Viral, Autoimmune, Hematologic and Bacterial Diseases (IMVAHB), IDMIT Department / IBFJ, 92265 Fontenay-aux-Roses, France; INSERM U1016, CNRS UMR8104, Université Paris Cité Institut Cochin, Paris, France; MIVEGEC, University of Montpellier, CNRS, IRD, Montpellier, France; Department of Bioengineering, Indian Institute of Science, Bengaluru, India

## Abstract

While most individuals suffer progressive disease following HIV infection, a small fraction spontaneously controls the infection. Although CD8 T-cells have been implicated in this natural control, their mechanistic roles are yet to be established. Here, we combined mathematical modeling and analysis of data from 16 SIV-infected macaques, of which 12 were natural controllers, to elucidate the role of CD8 T-cells in natural control. For each macaque, we considered, in addition to the canonical *in vivo* plasma viral load and SIV DNA data, longitudinal *ex vivo* measurements of the virus suppressive capacity of CD8 T-cells. Available mathematical models do not allow analysis of such combined *in vivo*-*ex vivo* datasets. By explicitly modeling the *ex vivo* assay and integrating it with *in vivo* dynamics, we developed a new framework that enabled the analysis. Our model fit the data well and estimated that the recruitment rate and/or maximal killing rate of CD8 T-cells was up to 2-fold higher in controllers than non-controllers (p=0.013). Importantly, the cumulative suppressive capacity of CD8 T-cells over the first 4-6 weeks of infection was associated with virus control (Spearman’s ρ=- 0.51; p=0.05). Thus, our analysis identified the early cumulative suppressive capacity of CD8 T-cells as a predictor of natural control. Furthermore, simulating a large virtual population, our model quantified the minimum capacity of this early CD8 T-cell response necessary for long-term control. Our study presents new, quantitative insights into the role of CD8 T-cells in the natural control of HIV infection and has implications for remission strategies.

## INTRODUCTION

Antiretroviral therapy (ART) suppresses viremia in individuals with HIV and arrests progression to AIDS but does not eradicate the virus^1^. Stopping treatment even after years of HIV control under ART typically results in viral recrudescence and disease progression. ART must therefore be administered lifelong. Enormous efforts are underway to devise interventions that could elicit long-term virus control following short-term drug exposure^2, 3, 4, 5^. These efforts are inspired by the rare individuals, termed ‘natural controllers,’ who control viremia without any intervention^6^.

Efforts to identify the determinants of natural control, in humans and non-human primates, point to the crucial role of CD8 T-cells in establishing such control. Natural controllers have an over-representation of the protective major histocompatibility complex (MHC) class-I haplotypes, like B*57 and B*27, which appear to facilitate strong, cross-reactive CD8 T-cell responses to HIV^7, 8, 9^. Natural controllers tend to have a higher frequency of polyfunctional^9, 10^ and Gag-specific^11, 12^ CD8 T-cells and suffer lower levels of CD8 T-cell exhaustion^13^ than non-controllers. Furthermore, memory-like CD8 T-cells were reported to develop early after infection in controllers^14^, which may confer protective immunity. Conversely, suboptimal CD8 T-cell responses were correlated with impaired virus control^13, 15, 16^.

Despite this substantial evidence, the processes determining CD8 T-cell response kinetics that underlie natural control are yet to be clearly elucidated. This is possibly because most studies offer either a static snapshot or a qualitative measure of the CD8 T-cell response, whereas the CD8 T-cell response is dynamic and influences disease outcome by its quality as well as magnitude^17^. Indeed, the frequency of the CD8 T-cells alone was found not to be a reliable indicator of natural control^10, 14, 18^.

In an effort to characterize the CD8 T-cell response more comprehensively, an *ex vivo* assay was developed some years ago^19^ and has since been employed in multiple studies on HIV and SIV infections^9, 11, 14, 20, 21, 22, 23, 24^. The assay measures the capacity of the CD8 T-cells drawn from an individual to suppress the viral load in a culture of autologous target CD4 T-cells exposed to the virus. This ‘suppressive capacity’ is thus a composite measure of the quality and the quantity of the CD8 T-cells. Furthermore, longitudinal measurements of the suppressive capacity provide a dynamic measure of the CD8 T-cell response during infection and hold promise of elucidating its mechanistic underpinnings in natural control. Because of the complex, nonlinear interactions between CD8 T-cells and antigen, however, identifying characteristics of the CD8 T-cell response associated with virus control would require analysis of the suppressive capacity measurements simultaneously with measurements of plasma viral load and other markers of disease state, such as the frequency of infected cells, using a mathematical model. Available mathematical models of virus dynamics have yielded profound insights into long-term HIV/SIV control^8, 25, 26, 27^ but are incapable of this analysis. The challenge arises from the multiscale and combined *in vivo-ex vivo* nature of the dataset, which current models cannot handle. Here, we developed a new mathematical model that enables this analysis. We made conceptual advances based on which our model not only described the suppressive capacity measurements but also explicitly incorporated the influence of the suppressive capacity on *in vivo* virus dynamics. We applied the model to analyze published data from an SIV-cynomolgus macaque model^14^, which showed robust maturation of virus-specific CD8 T-cell responses in natural controllers. We found that the cumulative CD8 T-cell suppressive capacity early in the infection was a correlate of natural control at later stages.

## RESULTS

### Model integrating *ex vivo* CD8 T-cell suppressive capacity with *in vivo* virus dynamics

We developed our model in three stages (Methods): First, we modeled virus dynamics in the *ex vivo* cultures, quantifying the CD8 T-cell suppressive capacity (Fig. S1). Second, we derived analytical expressions linking the suppressive capacity with the killing rate of infected cells by CD8 T-cells. Third, we incorporated the killing rate into models of *in vivo* virus dynamics, thereby constructing a unified model capable of predicting the measured *in vivo* and *ex vivo* quantities. We tested variants of the *in vivo* model using a formal model building strategy (Text S1, Fig. S2 – S6, Table S1 – S6) to identify the best model (Methods). The following equations describe the resulting model (Fig. 1):

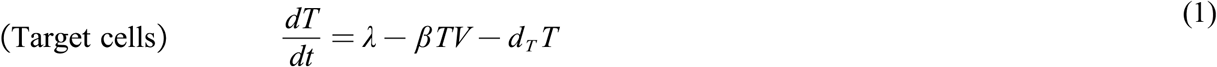

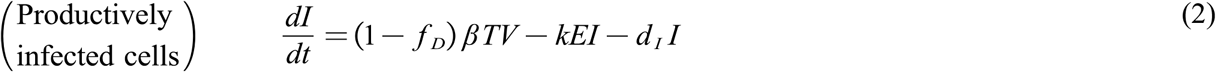

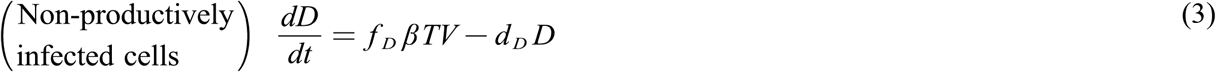

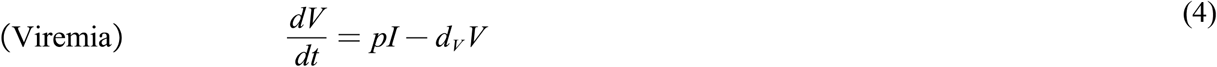

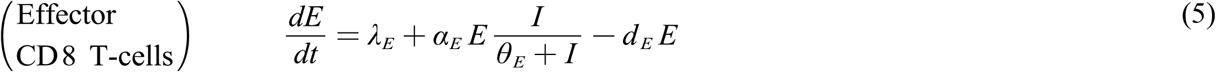

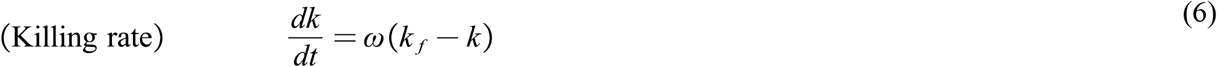

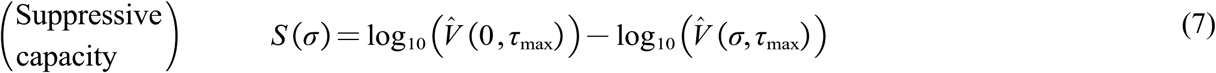

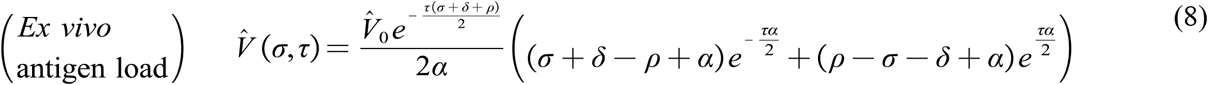

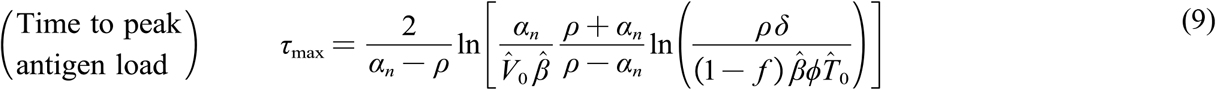

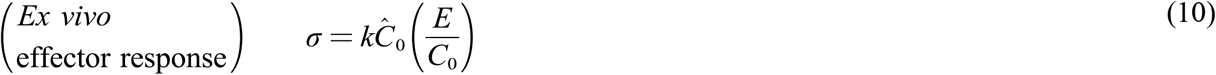

where and 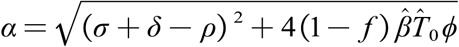.

**Fig. 1:**
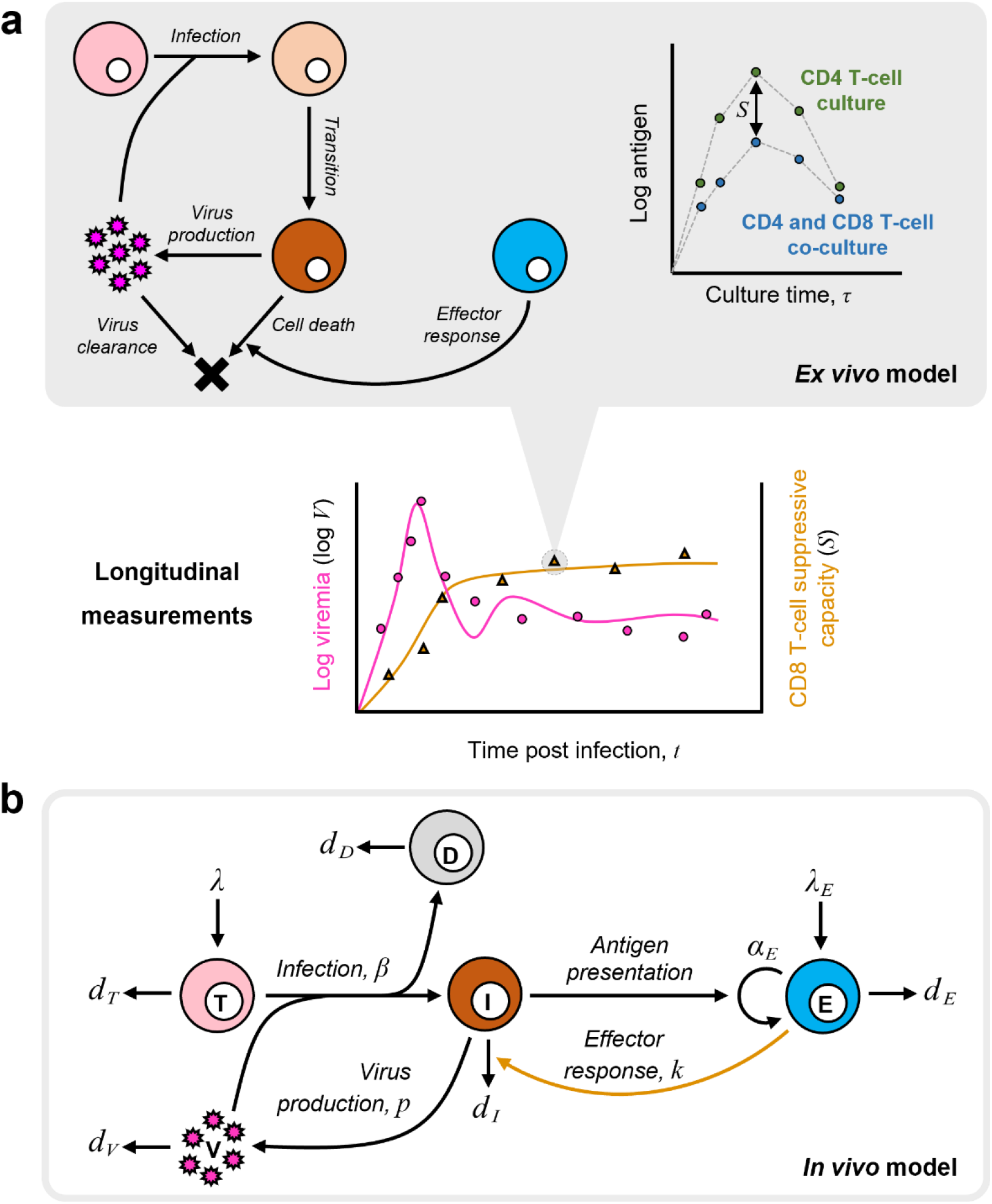
Schematic of the mathematical model. **(a) Model of the *ex vivo* assay**. The events in the *ex vivo* cultures (left) leading to the dynamics (right) and the reported suppressive capacity (***S***) as the difference in the antigen load in the cultures with and without CD8 T-cells. The model enables prediction of ***S*** and hence analysis of its longitudinal measurements along with *in vivo* measurements such as viremia (bottom), when integrated in a model of *in vivo* dynamics. **(b) Model of *in vivo* dynamics**. The events driving *in vivo* infection contained in our model, including the CD8 T-cell suppressive capacity reflected in the effector response (yellow arrow), linking the *ex vivo* and *in vivo* datasets (Methods).

Here, uninfected CD4 T-cells, *T*, are recruited at the rate *λ* and die at the rate *d*_*T*_*T*. They get infected by free virions in plasma, *V*, at the rate *βTV*. Because infected cell numbers are typically proportional to the viral load (see below), the latter infection rate subsumes cell-cell transmission^28, 29^. A fraction *f*_*D*_ of these infections results in non-productively infected cells, *D*, which do not produce virions. The remaining fraction, 1 − *f*_*D*_, results in productively infected cells, *I*, which produce virions at the rate _*p*_*I* . The productively infected cells die due to virus-induced cytopathicity at the rate *d*_*I*_*I* or due to killing by virus-specific CD8 T-cells, *E*, at the rate *kEI*. Free virions are cleared at the rate *d*_*v*_*V*. The cells are produced at the rate *λ*_E_ and die at the rate *d*_*E*_*E*. They proliferate with the rate constant *α*_*E*_ and display a saturating dependence on the antigen level for activation, with the *θ*_*E*_ half-maximal saturation constant. The killing rate constant, *k*, depends on the quality of the effector response. For a given effector population *E*, a more focused effector response would imply a higher *k* . *k*. can thus vary with time due to clonal expansion, memory recall, exhaustion, and/or viral evolution^14, 30, 31, 32^. Here, we focused on untreated infection, where the quality of CD8 T-cells is expected to rise during early infection and eventually plateau^33, 34, 35, 36, 37^. We developed an empirical equation to capture this expected evolution of *k*. Accordingly, *k* rises from zero exponentially and saturates at *k*_*f*_, with the changes occurring over the timescale 1/ *ω*. CD8 T-cells can also have non-cytolytic effects on infected cells^38^. Here, we focused on their cytolytic effects, which were found to be dominant in the *ex vivo* assays^9, 14^.

*S* is the suppressive capacity measured using the *ex vivo* assay. In the experiments, it is estimated as the difference between the antigen load in the CD4 T-cell cultures exposed to the virus in the absence, 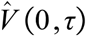, and presence, 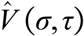, of CD8 T-cells, measured at the time τ_max_ when the antigen load peaks in the former culture^19^. At any time *t* during the *in vivo* infection, *S* is estimated based on the CD4 and CD8 T-cells drawn from the infected macaque at the time *t* for the *ex vivo* assays. *S* is determined to be a function of *σ*, the elimination rate of infected cells in culture due to CD8 T-cells. *σ* is thus the product of the killing rate constant and *k* the population of CD8 T-cells employed in the assay that are virus-specific. 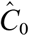 is the total population of CD8 T-cells in the assay, of which the fraction *E*/*C*_0_ is virus-specific, where *C*_0_ is the total CD8 T-cell concentration in the host. *σ* thus links the *ex vivo* observations with the *in vivo* dynamics. The other parameters in the expressions for 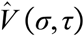 and τ_max_ are associated with the *ex vivo* assay (Table S7) and are described in *S* 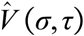 the Methods along with a detailed derivation of the expressions for, (Methods) and (Text S2).

The above model offered the unified framework necessary for the simultaneous analysis of longitudinal *in vivo* measures of viral dynamics and *ex vivo* CD8 T-cell suppressive capacity. We applied the model to the analysis of data from SIV-infected macaques.

### Model recapitulated dynamics of all the markers

We considered longitudinal data of plasma viremia, SIV DNA levels and CD8 T-cell suppressive capacity from 16 cynomolgus macaques infected with SIV (Methods). We fit our model to the data using a nonlinear mixed effects approach (Methods). Our model provided excellent fits to the data (Fig. 2; Fig. S6). The estimated population parameters for the best-fit model are in Table 1, and the individual macaque parameters are in Table S6. The parameter estimates were consistent with previous reports, where available (see Discussion). All the measurements, *in vivo* and *ex vivo*, were thus recapitulated by our model.

**Table 1:** Population parameter estimates for the best-fit model. Estimates of the parameters from fitting the best-fit model (model #1, Table S1) to the macaque data (Fig. 2). Standard errors are in parentheses. *d*_*I*_, *θ*_*E*_, *d*_*E*_ and were fixed based on previous studies (Methods).

| Parameter (Units) | Fixed effect | Random effect |
| --- | --- | --- |
| $\lambda$ (cells mL<sup>-1</sup> d<sup>-1</sup>) | 352.70 (125.87) | 1.32 (0.26) |
| $\log_{10} \beta'$ (log mL cells<sup>-1</sup> d<sup>-1</sup>) | -2.84 (0.02) | - |
| $f_D$ | 0.93 (0.01) | 0.17 (0.03) |
| $\log_{10} \lambda_E^*$ (log d<sup>-2</sup>) | 0.15 (0.01) | 0.25 (0.11) |
| $d_I$ (d<sup>-1</sup>) | 0.10 | - |
| $d_D$ (d<sup>-1</sup>) | $6.9 \times 10^{-2}$ ($1.9 \times 10^{-2}$) | 0.69 (0.18) |
| $\gamma$ (cells<sup>-1</sup>) | 427.65 (110) | 0.66 (0.21) |
| $\alpha_E$ (d<sup>-1</sup>) | 0.63 (0.09) | 0.20 (0.15) |
| $\theta_E$ (cells mL<sup>-1</sup>) | 0.10 | - |
| $d_E$ (d<sup>-1</sup>) | 1.00 | - |
| $\log_{10} \omega$ (log d<sup>-1</sup>) | -2.50 (0.16) | 0.47 (0.11) |
| $\log_{10} T(0)$ (log cells mL<sup>-1</sup>) | 4.21 ($1.60 \times 10^{-2}$) | - |

**Fig. 2:**
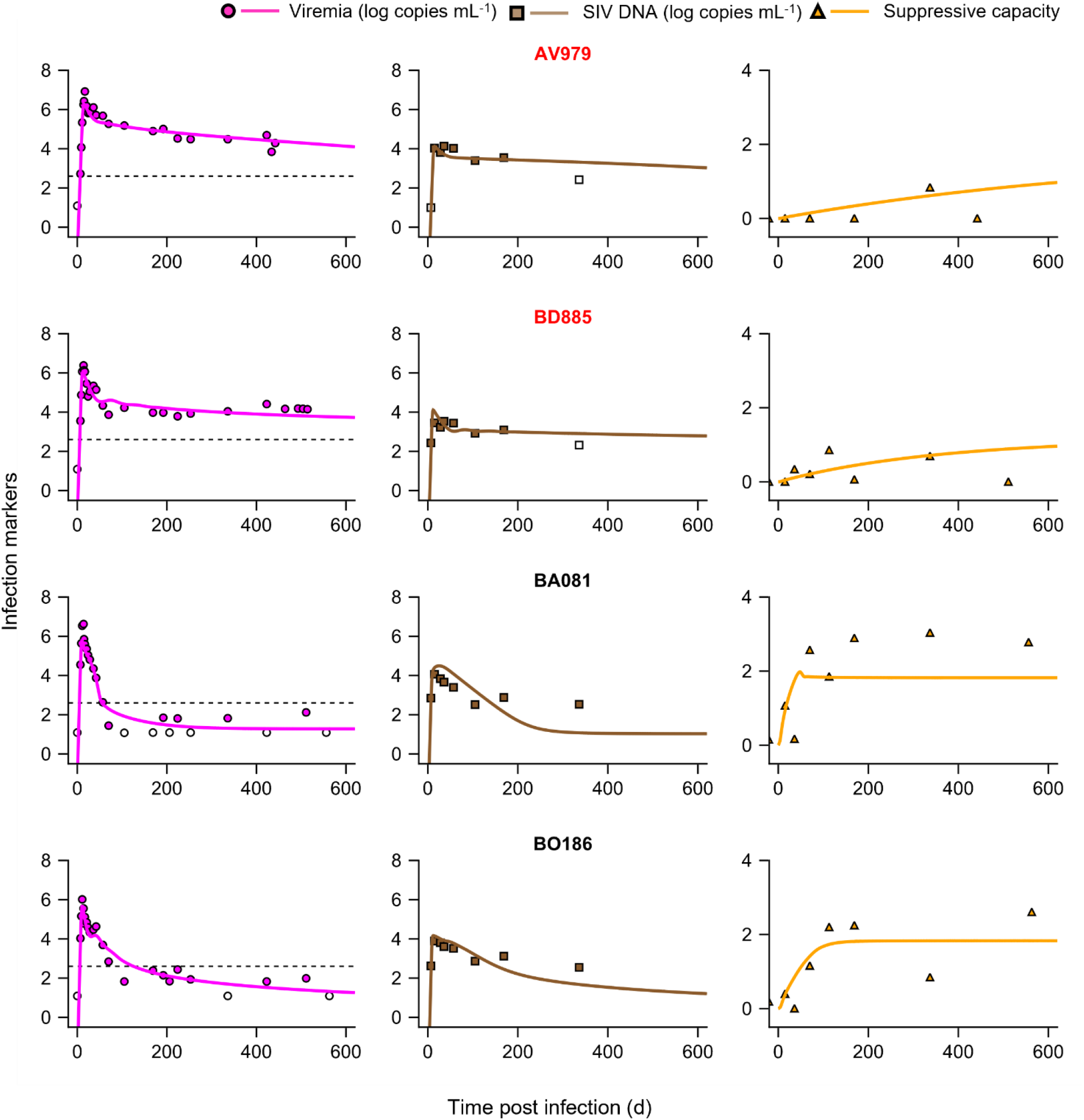
Model fits longitudinal *in vivo* virological and *ex vivo* suppressive capacity data. Model predictions (lines) from simultaneous fitting of the best-fit model (Methods) to all the three datasets (symbols), namely, viremia (left panels), SIV DNA (middle panels) and suppressive capacity (right panels). Macaques highlighted in red were progressors while those in black were controllers. The dashed line in the left panels indicates 400 copies mL^-1^. Open symbols are below the limit of detection. The predictions for the remaining 12 macaques are presented in Fig. S6. The resulting population parameter estimates are in Table 1 and individual parameter estimates are in Table S6.

Controllers in the experiment were identified as macaques that brought the viral load below 400 copies mL^-1^ after the primary infection phase and maintained it below this limit throughout (ref ^14^; Methods). By this definition, the dataset had 12 controllers and 4 progressors (or non-controllers). Our model fits yielded set-point viral loads above 400 copies mL^-1^ in the four progressors and below 400 copies mL^-1^ in all controllers, consistent with the experimental observations. Sensitivity analysis showed that these predictions were robust to parameter variations (Fig. S7). Using our best-fit model and parameter estimates, we assessed next the differences between controllers and progressors, focusing on CD8 T-cell responses.

### CD8 T-cell responses had greater antiviral capacity in controllers than progressors

Comparing best-fit parameter estimates, we found that controllers had a significantly higher recruitment rate and/or maximal killing rate of CD8 T-cells, contained in the composite parameter 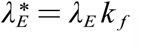, than progressors (Fig. 3a). Specifically, the median value of 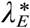 was 1.65 d^-2^ in controllers and 0.86 d^-2^ in progressors, implying a nearly 2-fold enhancement in controllers (p=0.013). Controllers also had a higher antigen-induced proliferation rate of CD8 T-cells (α_*E*_), although the latter difference was not significant (Fig. 3b). Thus, the CD8 T-cell response seemed more robust in controllers. The controllers, also, interestingly, had a lower value of the ratio of viral production and clearance rates, *γ* (Fig. S8), possibly due to innate immune responses or other cytokine-mediated effects which curtail viral production^14^. The other parameters were not significantly different between the groups (Fig. S8). Here, our aim was to assess whether CD8 T-cell responses would yield a correlate of natural control, notwithstanding other factors. We therefore focused on the differences in 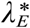 and α_*E*_, which would manifest as a difference in the suppressive capacity of the CD8 T-cells. Using the best-fit parameter estimates, we predicted the early time-course of the suppressive capacity, *S*, for all the macaques and found that the predicted *S* was significantly higher at day 28 in the controllers than progressors (Fig. 3c). This suggested that the early suppressive capacity of the CD8 T-cells could be a predictor of natural control. We evaluated this possibility next.

**Fig. 3:**
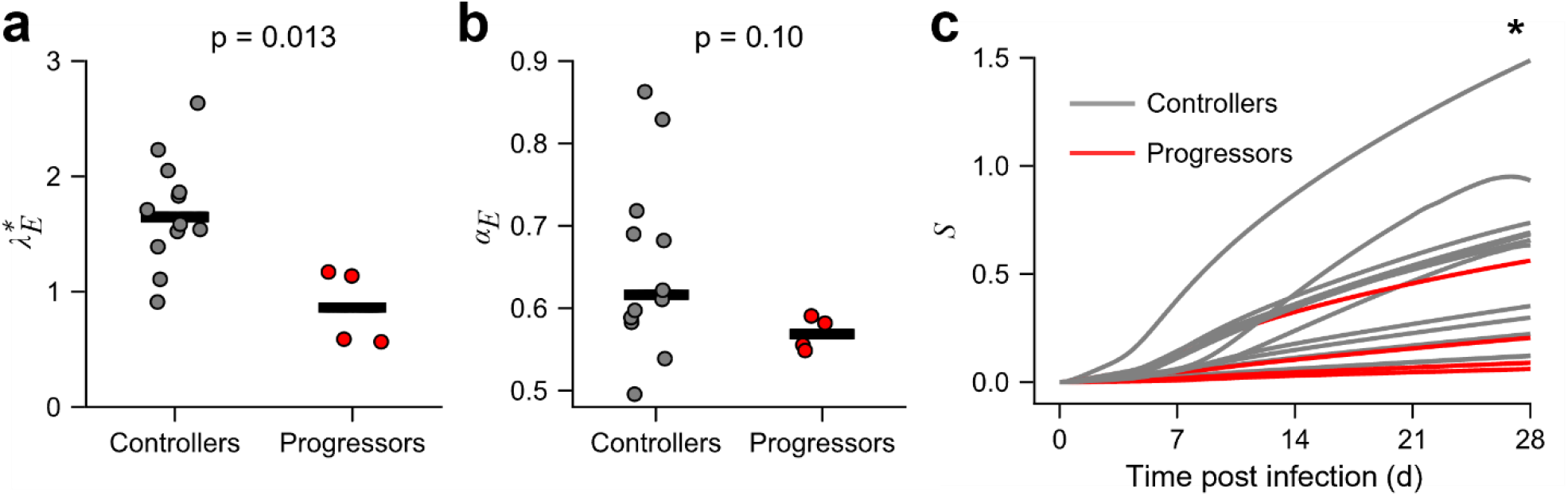
Natural controllers elicit stronger CD8 T-cell responses than progressors. Best-fit model predictions (Fig. 2) showed a higher **(a)** recruitment/killing rate and **(b)** antigen-induced proliferation rate of CD8 T-cells in controllers (gray) compared to non-controllers (red). Each symbol represents a macaque and the bar is the median. **(c)** Predictions using the best-fit parameters showed higher suppressive capacity in controllers than non-controllers. * indicates p=0.04 at the last time point using a Mann-Whitney U test.

### Cumulative antiviral capacity of the early CD8 T-cell response was correlated with viral control

By sampling parameter values from the distributions obtained in our fits above (Methods, Table S6), we generated a virtual population of 10^5^ macaques and simulated the progression of SIV infection in each using our model (Fig. 4a, 4b). We found that the range of set-point viral loads realized (10^0^ – 10^6^ copies mL^-1^) was consistent with the range observed in individuals with HIV^39^. For each virtual macaque, we computed the time-averaged area-under-the-curve of *S* over the first 28 days of infection, which we denoted *S*_*28*_. We found, interestingly, that *S*_*28*_ was inversely correlated with the set-point viral load (Fig. 4c). Thus, early CD8 T-cell responses with greater antiviral capacity were associated with lower set-point viral loads. Using data of the set-point viral loads from the 16 macaques above and the corresponding best-fit predictions of *S*, we found that the above correlation held also in the macaques we studied (Fig. 4d). Thus, the cumulative antiviral capacity of the early CD8 T-cell response was a predictor of viremic control.

**Fig. 4:**
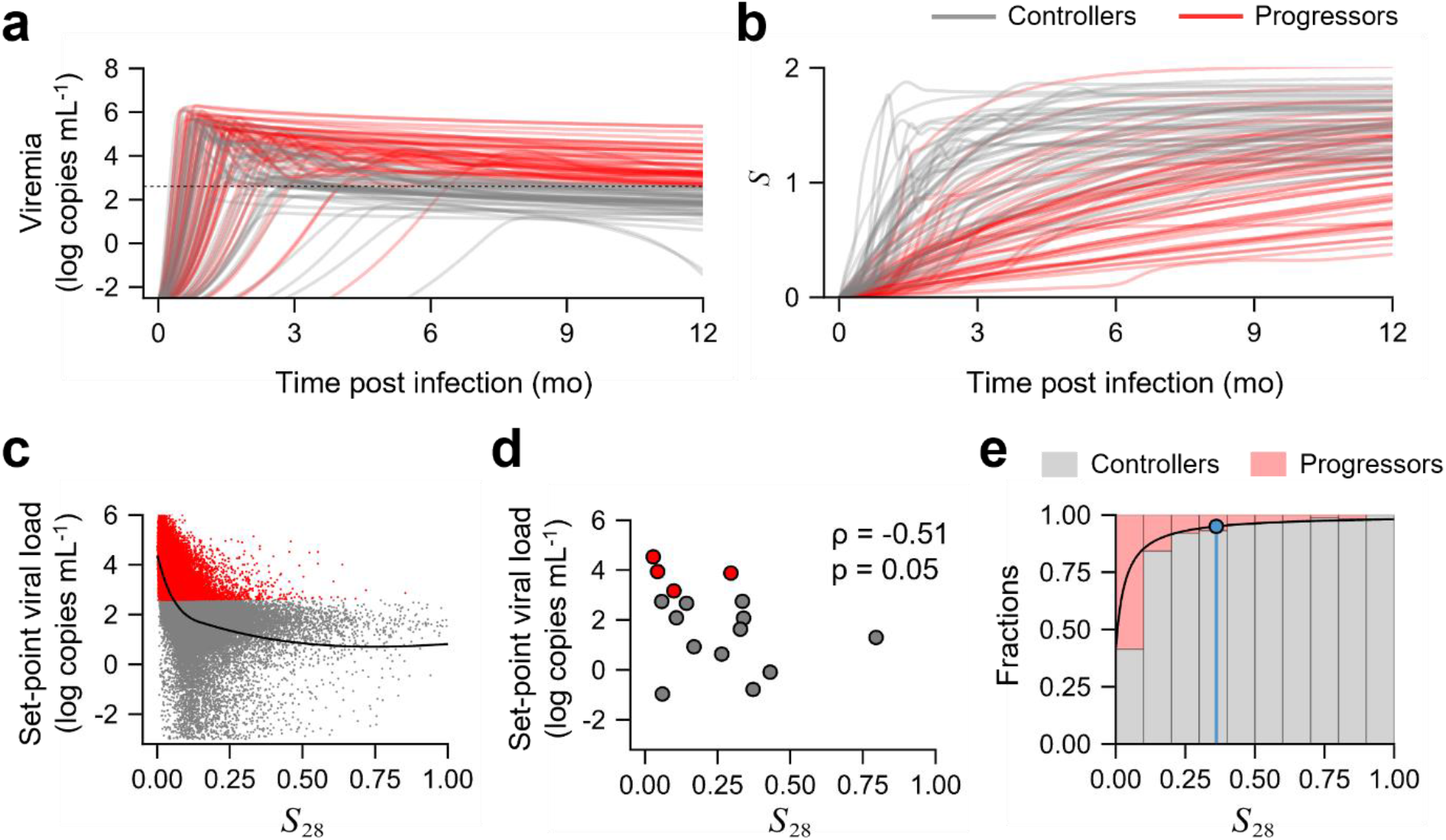
Early cumulative suppressive capacity is a marker of natural control. Dynamics of **(a)** viremia and **(b)** suppressive capacity predicted for virtual patients using our best-fit model. Trajectories for fifty controllers and fifty progressors are shown. Black dashed line indicates 400 copies mL^-1^. Correlation between set-point viral load and cumulative suppressive capacity *S*_28_ (see text) for **(c)** 100000 simulated individuals and **(d)** the 16 macaques studied. The black curve in **(c)** is a LOESS regression curve to visualize the inverse correlation. **(e)** The fraction of virtual individuals achieving control (gray bars) or experiencing progressive disease (red bars) as a function of *S*_28_. Each bar has of width 0.1 units of *S*_*28*_. The black curve is a fit of the estimated fractions to a first-order Hill function (Methods). The blue line represents the minimum *S*_28_ for >95% controllers, with control defined as set-point viral load <400 copies mL^-1^. Spearman’s ρ was calculated for assessing the correlations.

A model that did not incorporate the suppressive capacity measurements (Table S1, Table S8) could fit viral load data well (Fig. S9, S10), as is the case with available models^26, 27, 40^, but could not distinguish between the CD8 T-cell responses in controllers and progressors (Fig. S11a) and, therefore, could not identify the above correlate (Fig. S11b).

The correlate was robust to the duration (28 d) for evaluating the early CD8 T-cell response. For instance, the correlate held when 42 d was used instead of 28 d; the correlation between *S*_28_ and set-point viral load was as strong as the correlation with *S*_28_ (Fig. S12a). The correlation was expectedly lost when the time period was too short or long. When the period was too short (14 d, *S*_28_), a significant CD8 T-cell response was yet to be mounted, whereas when it was too long (90 d, *S*_*90*_), the early dynamics was masked by the dynamics in the chronic phase (Fig. S12a).

We asked next whether a threshold *S*_*28*_ existed that was associated with the set-point viral load of 400 copies mL^-1^ and could thus facilitate distinguishing controllers from progressors as defined in the experiments^14^. We found from the above virtual population that as *S*_*28*_ increased, the fraction of macaques that exhibited control increased (Fig. 4e). The fraction was ∼40% when *S*_*28*_ was <0.1 and rose to ∼95% when *S*_*28*_ was ∼0.4. Thus, we defined 0.4 as the critical *S*_*28*_, above which the chance of achieving viremic control was >95% in our predictions. We recognize that the threshold *S*_*28*_ would depend on the level of viremia used to define control; a more stringent definition (set-point viremia lower than 400 copies mL^-1^) would lead to a higher threshold (Fig. S12b). Nonetheless, once the set-point viremia for control is defined, the corresponding threshold *S*28 identified by our model offers a novel, measurable, early predictor of natural control. Indeed, with the 16 macaques we studied, the proportion of macaques exhibiting control increased with *S*28, consistent with our predictions (Fig. 4e). Further, although the sample size was small, macaques with *S*28 values above the predicted threshold of 0.4 were all found to exhibit control.

## DISCUSSION

Identifying correlates of natural control of HIV infection has been a long-standing goal. Here, combining mathematical modeling and analysis of longitudinal *in vivo* and *ex vivo* data from SIV-infected cynomolgus macaques, we identified the cumulative response of CD8 T-cells during the first 4-6 weeks of infection as an early, measurable marker of natural control. The more efficient was the early CD8 T-cell response, measured in terms of its cumulative virus suppressive capacity, the lower was the set-point viral load. The marker was robust to the duration (∼4-6 weeks) over which the early CD8 T-cell response was measured. To our knowledge, this is the first study to identify a quantitative marker predictive of long-term natural control without antiretroviral treatment.

We made significant advances in mathematical modeling that enabled the identification of the marker. The model had to contend with data that was a combination of *in vivo* virological and *ex vivo* immunological measurements. Furthermore, the measurements involved nested time courses. Specifically, each CD8 T-cell suppressive capacity measurement was obtained from time course data of antigen load from *ex vivo* cultures. Longitudinal measurements of suppressive capacity during the *in vivo* infection thus had *ex vivo* assay time course datasets nested within each measurement. Mathematical models thus far have not analyzed such combined *in vivo-ex vivo* datasets. Besides, standard fitting algorithms cannot routinely handle nested time course datasets. By analyzing the *ex vivo* assays, we developed an analytical expression that yielded the suppressive capacity as a function of the killing rate of infected cells by CD8 T-cells. This eliminated the need to consider the *ex vivo* time courses. It also allowed us to link the suppressive capacity measurements with *in vivo* infection dynamics. Consequently, like the plasma viral load, the CD8 T-cell suppressive capacity became a quantity that could be predicted by our model, and therefore fit simultaneously with longitudinal measurements of viral load and SIV DNA using standard fitting algorithms. Compared to available models^8, 26, 27, 40^, which typically rely on viral load and SIV DNA measurements alone, an extra dimension of information by way of the CD8 T-cell suppressive capacity measurements thus became accessible for constraining our model. This allowed more accurate inferences of the *in vivo* dynamics and, in particular, enabled identification of the above marker of natural control, which available models missed.

The quality of the fits (Fig. 2) as well as the consistency of the best-fit parameter values with independent estimates, where available, gave us further confidence in our inferences. For instance, the best-fit value of the fraction of infection events resulting in non-productively infected cells (*f*_*D*_) was 0.93, close to independent estimates of 95% infection events turning abortive^40, 41, 42^. The best-fit initial target cell concentration was ∼16 cells μL^-1^, which corresponds to ∼3% of the baseline CD4 T-cell count in blood (median 654 cells μL^-1^; ref ^14^), again consistent with ∼5% of the CD4 T-cells in blood expressing CCR5^43^, required for SIVmac251 infection. The best-fit ratio of viral production and clearance rates, *γ*, was ∼400, consistent with previous reports^26, 44, 45^. The best-fit timescale of the evolution of the quality of the CD8 T-cell response (In 2/ ω) was ∼220 days. While the processes driving this timescale are yet to be established, it was comparable to the timescale of evolution of viral diversity (months to years)^30, 31^.

Our study offers new insights into the potential role of CD8 T-cells in establishing natural control of HIV infection. While several studies have measured CD8 T-cell responses during infection, including in its early stages, the measurements have proven inadequate to distinguish between controllers and progressors^14, 18^. Thus, despite the recognition of the importance of CD8 T-cells, a major gap existed in our understanding of their specific role in natural control. Our study makes an important advance by accounting more comprehensively for the antiviral activity of CD8 T-cells than has been done thus far in describing virus dynamics. Our formalism considered not only the quality and the quantity of the CD8 T-cell response but also its time course during the infection. The cumulative antiviral capacity of the CD8 T-cell response early (∼4-6 weeks) in infection thus emerged as a correlate of long-term virus control. In our analysis, the higher antiviral capacities in controllers were attributable to greater recruitment rates and/or maximal killing rates of CD8 T-cells compared to progressors. Future studies may assess further the specific implications of this early cumulative response, such as the restriction of the latent reservoir^46, 47^, the prevention or reversal of CD8 T cell exhaustion^25, 26, 46^, and/or the formation of an adequate memory pool^14^, that may underlie the long-term control realized.

We anticipate implications of our findings for the ongoing efforts to elicit long-term HIV remission^4^. First, using the distribution of parameter values based on fits of our model to the macaque data, our study identified a threshold strength of the marker (*S*_28_) for achieving a set-point viremia representative of long-term control. Future studies may translate this threshold to humans, thereby predicting quantitative targets for interventions aimed at eliciting potent early CD8 T-cell responses for achieving lasting control of HIV-1 infection. Such interventions include vaccination strategies^17^ as well as immunotherapies with immune checkpoint inhibitors^48^ and broadly-neutralizing antibodies^49^ aimed at eliciting better CD8 T-cell responses. Second, early interventions with ART have been shown to increase the chances of achieving post-treatment control^32, 50^. While CD8 T-cell responses have been implicated in the establishment of such control^25, 26, 32, 50^, how ART may trigger such responses is unclear. Our study suggests that supplementing measurements of viral load with those of *ex vivo* CD8 T-cell suppressive capacity may help elucidate the underlying mechanisms. Such data could be analyzed using the modeling framework developed in our study. The analysis may help understand whether natural control and post-treatment control are the same state realized via two different routes or are two fundamentally distinct states. If the correlation between CD8 T-cell responses early after treatment cessation and the ensuing set-point viral load were similar to that observed in the present study, then post-treatment control would likely be the same state as natural control. Then the notion of the threshold CD8 T-cell response we identified here may be translated also to the post-treatment control scenario. This would further inform the many strategies being explored today that combine ART with other interventions, the latter often designed to improve CD8 T-cell responses, to achieve long-term control^4, 17, 51^.

Our study has limitations. First, although our model is complex, it considered the most parsimonious description of *in vivo* dynamics based on the data available. It thus did not include processes like CD8 T-cell exhaustion or memory. While the model successfully recapitulated the datasets from the untreated macaques that we examined, extending it to treated macaques may require explicitly considering the latter processes. Such advances may also alter the dynamical features of the model–for instance, by introducing bistability^25, 26, 52^–the implications of which remain to be ascertained. Second, we employed an empirical framework to describe the evolution of the quality of the CD8 T-cell response with time. Again, while such a framework may be adequate for recapitulating natural control, a mechanistic framework, involving phenomena such as CD8 T-cell clonal expansion, differentiation to memory phenotypes and recall^14, 53^, may be required in other scenarios. Third, the data used in this study was from a non-human primate cohort that had an unrealistically large percentage of natural controllers compared to what is observed in humans^7, 54^. In our virtual populations, we ascertained that the range of set-point viral loads predicted by our model was consistent with humans. Yet, to translate the threshold value of the cumulative suppressive capacity to humans, parameters recapitulating not only the range but also the distribution of set-point viremia in humans would have to be employed. Fourth, our model assumed that killing of infected cells was the dominant mode of CD8 T-cell effector function. This is based on observations of the loss of suppressive capacity when contact between CD8 T-cells and infected cells was eliminated in the *ex vivo* assays^9, 14^. Nonetheless, non-cytolytic effects of CD8 T-cells have been observed *in vivo*^38, 55, 56^. Future studies may incorporate both cytolytic and non-cytolytic effects and refine our predictions based on their relative contributions^38^.

In summary, our study identified a new, robust early marker of natural control of HIV infection, which not only advances our understanding of the mechanisms driving such control but also informs ongoing efforts to devise strategies for eliciting lasting HIV remission.

## METHODS

### Model development

#### *Model of* ex vivo *virus dynamics*

We first considered the *ex vivo* assay of the CD8 T-cell suppressive capacity measurements. Here, a fixed number of target CD4 T-cells drawn from an individual is exposed to free virions in culture either in the absence or the presence of a fixed number of autologous CD8 T-cells drawn simultaneously and the time course of the antigen load in the supernatant is measured. We developed a model to predict the latter time course. The suppressive capacity is estimated as the extent to which the antigen load is reduced in the presence of CD8 T-cells compared to its peak level in their absence.

The following equations describe the virus dynamics in a culture of target CD4 T-cells exposed to free virions (Fig. 1a):

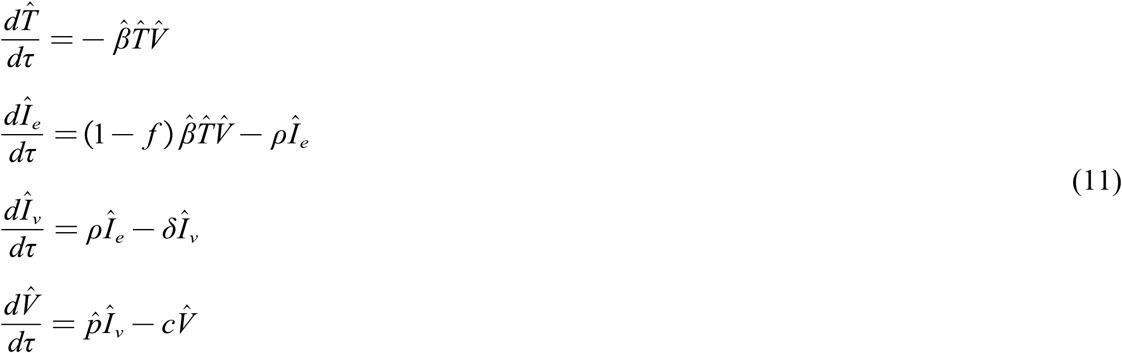

Here, target cells, 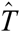, get infected by virus 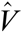 at the rate 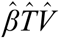. A fraction of these 1 − *f* infection events is productive, giving rise to infected cells in the eclipse phase, 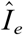, from which virus production is yet to occur. The remaining fraction of the infection events results in non-productively infected cells. These latter cells are assumed to be rendered non-susceptible because of subsequent CD4 downregulation^57, 58^. Infected cells in the eclipse phase transition to virus-producing cells, 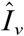, at the rate 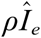, and produce free virions at the rate 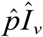. Free virions are cleared at the rate 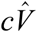. The elimination rate of virus-producing cells is 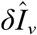 is the time from the start of the infection in culture. Further, we use the viral load, 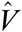, as a proxy for the antigen load, such as p24 (capsid protein) or p27 (non-structural regulatory protein) levels used as a marker of viral production^19^ in the assay.

Because virus production and clearance are fast compared to infection^28, 29^, we assumed a quasi-steady state for the dynamics of in 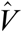 equation (11), so that 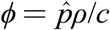 and 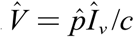. We fit the model to longitudinal data of antigen levels measured in *ex vivo* CD4 T-cell cultures. The model fit the data well (Fig. S1a), recapitulating the rise and fall of antigen with time, and yielded estimates of *ρ* (Table S7).

Next, to estimate the suppressive capacity, we applied the same model as above to data from co-cultures with a 1:1 mixture of CD4 T-cells and CD8 T-cells exposed to the virus. We let the elimination rate of virus-producing cells in equation (11) be 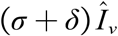, where *δ* is the death rate constant of infected cells due to virus-induced cytopathicity and *σ* is the increase in the death rate constant due to CD8 T-cells. Because the population of CD8 T-cells or their killing efficiency is not expected to change during the timeframe of the assays, σ can be assumed to be a constant. Our model fit the time-evolution of the antigen load in the co-culture assays (Fig. S1b) and yielded estimates of σ. From the fits, the difference in the antigen load at the time when the antigen level peaks in the CD4 T-cell monoculture can be calculated, linking σ to the reported suppressive capacity, *S*, of the CD8 T-cells.

The above procedure, however, estimates σ by analyzing the entire time-course of the monoculture and co-culture assays at any *in vivo* measurement time point (Fig. 1), which would render data fitting intractable. We therefore developed approximations that would yield an analytical expression linking σ and *S*.

#### *Linking* ex vivo *suppressive capacity to* in vivo *killing rate of CD8 T-cells*

Our approach was the following. First, we derived an analytical expression of the time-evolution of the antigen load in the *ex vivo* assay. Second, we obtained an analytical expression of the time at which the antigen load would attain its peak. With these two expressions, we predicted the difference in the antigen load between the monoculture and co-culture assays at the time when the load peaked in the monoculture, thus yielding *S* as a function of σ. We present details below.

We recognized that the target cell population remained close to its initial value nearly all the way until the peak in the infection in the monoculture (Fig. S1a). We therefore assumed that the target cell population was constant, i.e., 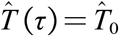, the initial target cell concentration, until the peak. This transformed our nonlinear model equations in (11) into the set of linear equations below:

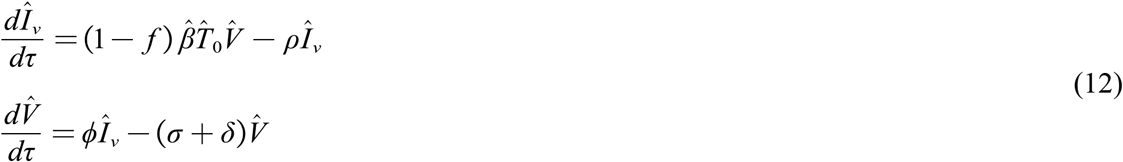

Solving equations in (12) for the viral load yielded

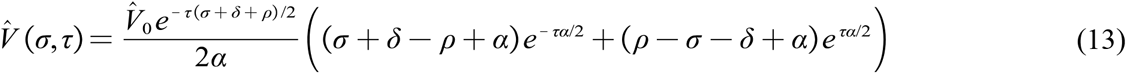

where 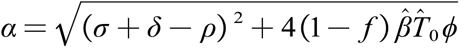. Predictions with this approximation (equation (13)) agreed well with the true solution (equation (11)) of the antigen load until the peak (Fig. S1c).

Next, we recognized, following epidemiological models^59, 60^, that the peak in the infection occurs when the effective reproductive ratio equals 1. Using next-generation matrix methods^59, 60^, we derived an analytical expression for the effective reproductive ratio (Text S2). This yielded τ_max_ as the time when the reduction in the target cell population due to the infection would drive the effective reproductive ratio to 1 (Text S2):

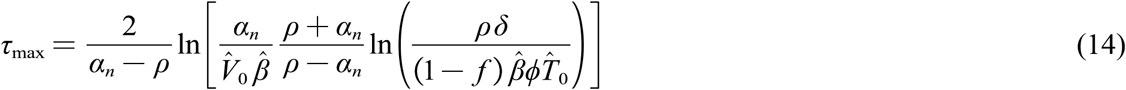

where 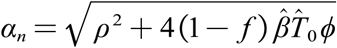. Combining the expressions of 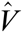 and 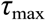 yielded the desired link between *S* and σ:

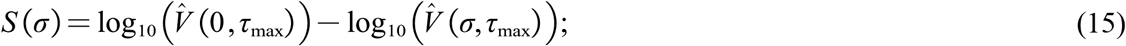

where 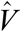 follows from equation (13).

Estimates of *S* (τ) obtained from equation (15) were close to those obtained by integrating equation (11) (Fig. S1d).

#### *Model building strategy for* in vivo *dynamics*

To identify the number of SIV DNA compartments our within-host models should contain, we fit mono-, bi-, and tri-exponential curves to the post-peak SIV DNA data. The SIV DNA data we used did not differentiate between unintegrated and integrated (intact and defective) SIV DNA. If *z* (*t*) represents the DNA level at time *t*, then a multi-exponential function is given by

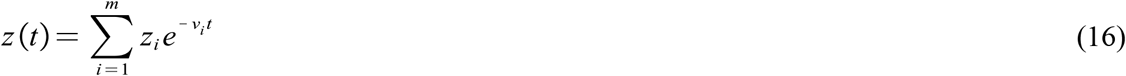

Here, *m* is the number of phases (or compartments), *v*_*i*_ represents the decay rate constant of the *i*^th^ phase, and *z*_*i*_ is the constant pre-factor for the *i*^th^ phase, respectively. The initial condition 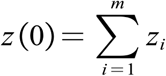 is the estimated DNA level in the blood at *t* = 0, the time point where the measurement peaked for the macaque. We found that a bi-exponential curve explained the data best (Fig. S13). Accordingly, we incorporated two SIV DNA compartments in our *in vivo* models.

Next, we constructed several models to describe the *in vivo* dynamics with two SIV DNA compartments (Text S1). We compared these models by fitting data. The best model, with the lowest Bayesian Information Criterion (BIC), is described in the Results. We analyzed the model for its structural identifiability and applied it to fit data.

### Structural identifiability of model parameters and data fitting

We analyzed the global structural identifiability of all models before fitting them to data using StructuralIdentifiability.jl package in Julia^61^. We had to fix the parameters *d*_*I*_, *θ*_*E*_, and *d*_*E*_, to make the remaining parameters of the best-fit model uniquely identifiable. Additional parameters had to be fixed in other tested models (Table S1), as they involved more parameters (Table S2 – S5).

From previous studies, we fixed *θ*_*E*_ = 0.1 cells mL^-1^ and *d*_*E*_ = 0.1 d^-1^ (ref ^25, 62^). We fixed *d*_*I*_ = 0.1 d^-1^ based on recent estimates of the half-life of productively infected cells (1.0 d to 1.7 d)^40, 56, 63^ and estimates of >40% of infected cell loss attributable to CD8 T-cell function^64^. Note that our model prediction of the set-point viral load was not sensitive to *d*_*I*_ (Fig. S7).

In the *in vivo* model, *k*_*f*_ was not identifiable. So, we applied the transformation *E*^*^ = *k*_*f*_ *E*. Also, viral production and clearance happen at a much faster rate than other *in vivo* processes^28, 29^. So, assuming quasi-steady state between virion production and clearance rates^28, 29^, we simplified the equation for viremia, giving us *pI* ≈ *d*_*V*_*V* ⟹ *V*(*t*) = *γ I*(*t*) where γ=*p/d*. These transformations to the *in vivo* model combined with the analytical expression linking *S* and *σ* for the *ex vivo* measurements yielded

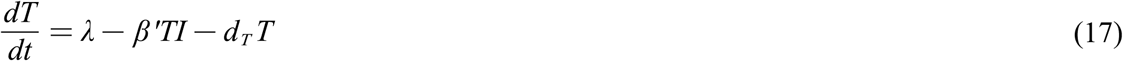

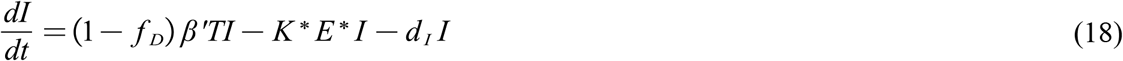

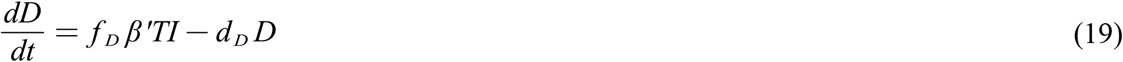

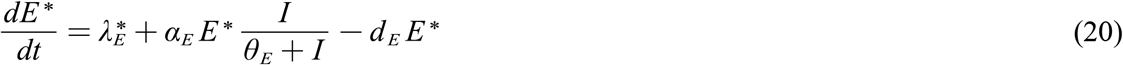

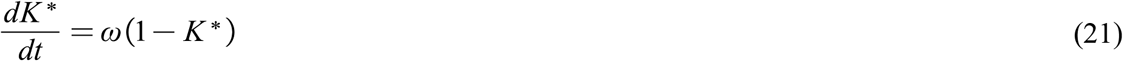

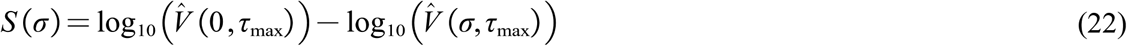

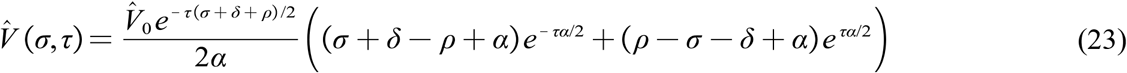

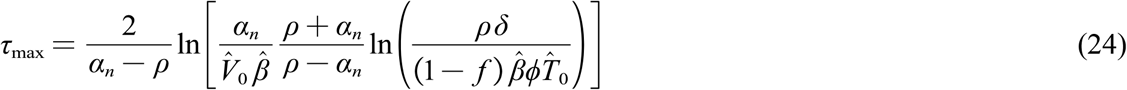

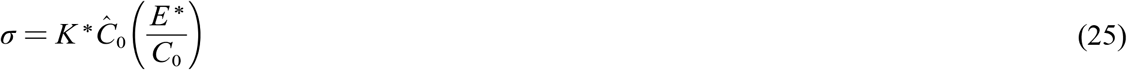

where 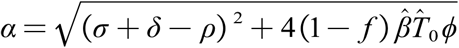 and 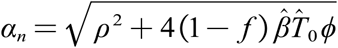. Here, β ′ = *γ′, K*^*^ = *k/k*_*f*_, *E*^*^ = *k*_*f*_ *E* and 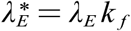. The above equations (17 - 25) were used for data fitting. Parameters used for the *ex vivo* model are presented in Table S7, and the initial conditions for the *in vivo* model are provided in Table S9.

### Statistical model for longitudinal data fitting

We employed the nonlinear mixed effects modeling (NLME) approach for fitting longitudinal data and used the implementation of stochastic approximation of expectation-maximization (SAEM) algorithm in Monolix 2021R1 (https://lixoft.com/). Initial conditions for the *in vivo* models are provided in Table S9. The variables *V* = _*γ*_*I, I* + *D* and *S* were fit to the viremia, SIV DNA, and suppressive capacity datasets, respectively. We assumed random effects for all parameters and removed them if they were less than 0.1. The statistical model describing these observations is

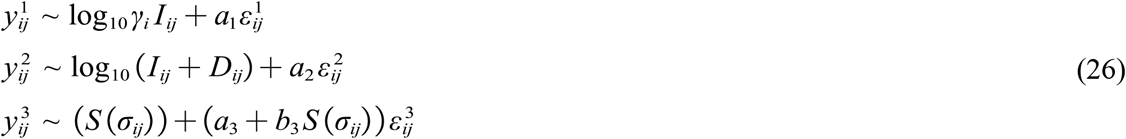

Here, *y*_*ij*_ represents the observations for the *i*^th^ individual at the *j*^th^ time point. The superscripts 1, 2 and 3 represent the log-transformed viremia, log-transformed SIV DNA and suppressive capacity measurements, respectively. is the residual Gaussian error with a constant standard deviation. Thus, for viremia and total SIV DNA datasets, we used a constant error model, while for the suppressive capacity data, both constant and proportional error terms were considered. Fits to the best-fit model are presented in Fig. 2 and Fig. S6, while for the other models, they are presented in Fig. S2 – S5.

### Sensitivity analysis

We performed sensitivity analysis of the set-point viral load estimates of our best-fit model. Sobol’s method was employed using the GlobalSensitivity.jl^61^ package in Julia.

### Virtual population

All parameters except for *f*_*D*_, which followed a logit-normal distribution with bounds between 0 and 1, were assumed to follow a log-normal distribution. Consequently, log_10_ *β* ′, 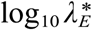, log_10_ *ω* and log_10_ *T*(0) followed a normal distribution. After model fitting, analytical forms of the corresponding distributions were fit to histograms of individual parameter values using a maximum likelihood estimation algorithm in Julia^65^. The virtual population (Fig. 4a – e) was then generated by sampling these parameters from the fit distributions.

The fraction of controllers estimated by our model, plotted in Fig. 4e, was fit to a first-order Hill function of *S*_28_ given by 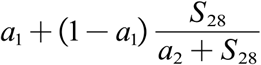 using the nonlinear Levenberg-Marquardt algorithm in Julia^65^. Here, *a*_1_ and *a*_2_ were fit parameters. Accordingly, *a*_1_ is the probability of achieving control in the limit of a negligible early CD8 T-cell response (*S*_28_ → 0) and is the half-maximal saturation constant.

### Data

We obtained data from a published study^14^. In the study, 16 macaques, of which 6 carried the protective M6 MHC haplotype, were infected with SIVmac251 intrarectally. They were then followed for 18 months without any intervention. Throughout this time, viremia, SIV DNA in blood and suppressive capacity of CD8 T-cells were measured at different time points. By the end of the study, 12 of the 16 macaques were identified as controllers. Viremia measurements were made as copies of SIV RNA mL^-1^ of blood. SIV DNA levels per million cells were converted from copies per 10^6^ leukocytes to copies mL^-1^ of blood, using individual blood leukocyte counts sampled simultaneously to the SIV DNA measurements.

## Codes and data availability

The codes employed for parameter estimation using NLME approach in Monolix and subsequent analyses using Julia 1.7.3^66^ are available on the GitHub repository https://github.com/vembha/SIV_natural_control. Spearman’s ρ was estimated using MATLAB 2023a Update 2.

## Acknowledgements

We thank Benoit Delache, Brice Targat, ClaireTorres, Christelle Cassan, Jean-Marie Robert, Julie Morin, Patricia Brochard, Sabrina Guenounou, Sebastien Langlois, and Virgile Monnet for expert technical assistance, Antonio Cosma for helpful discussion, Lev Stimmer for anatomopathology expertise, Delphine Desjardins and Isabelle Mangeot-Méderlé for helpful project management at IDMIT, and Christophe Joubert for veterinarian assistance at the animal facility at CEA. The SIV1C cell line was kindly provided by François Villinger. The SIVmac239 Gag Peptide Set was obtained through the NIH AIDS Reagent Program, Division of AIDS, NIAID, NIH. FTC, DTG and TDF were obtained from Gilead and ViiV healthcare though the “IAS Towards an HIV Cure” common Material Transfer Agreement for preclinical studies in HIV cure research. We thank Pranesh Padmanabhan and Rajat Desikan for their comments on the manuscript.

## Author contributions

B.V., A.S.-C., J.G., N.M.D., C.P. designed the study. C.P., A.M., V.A.-F., R.D.-D., N.D.-B., R.L.G., C.R., B.V., A.S.-C. performed the experiments and provided the data. J.G., B.V., A.S.-C., N.M.D. supervised the study. B.V., V.M. developed the models and codes and performed calculations. B.V., V.M., J.G., N.M.D. analyzed the data and interpreted the results. All authors wrote the paper and edited the paper.

