## Supplementary information for "Antiviral capacity of the early CD8 T-cell response is predictive of natural control of SIV infection"


**Supplementary materials:** Text notes: 2; Figures: 13; Tables: 9; References: 17

#### TEXT S1. *IN VIVO* MODEL VARIANTS

We describe variants of the *in vivo* model (model #1; Methods, main text; Table S1) we considered based on previous studies on HIV control[1, 2, 3, 4, 5]. First, following Conway and Perelson[1], we built model #2 (equation (S1)) with SIV-specific effector CD8 T-cells getting exhausted at the rate $\alpha_X E \frac{I}{\theta_X + I}$. The other equations in the model are identical to those in model #1.

$$
\begin{aligned}
\frac{dT}{dt} &= \lambda - \beta' TI - d_T T \\
\frac{dI}{dt} &= (1 - f_D)\beta' TI - K^* E^* I - d_I I \\
\frac{dD}{dt} &= f_D \beta' TI - d_D D \\
\frac{dE^*}{dt} &= \lambda_E^* + \alpha_E E^* \frac{I}{\theta_E + I} - (\alpha_E + \alpha_R) E^* \frac{I}{\theta_X + I} - d_E E^* \\
\frac{dK^*}{dt} &= \omega(1 - K^*)
\end{aligned}
\qquad \text{(S1)}
$$

Here, $\alpha_X$ is the exhaustion rate constant, and $\theta_X$ is the half-maximal saturation constant for the exhaustion rate dependent on the instantaneous antigen level. To ensure that the estimate of $\alpha_X$ is larger than $\alpha_E$[1, 2], we used $\alpha_X = \alpha_E + \alpha_R$ and constrained $\alpha_R$ to be strictly positive.

Model #3 tracks the exhaustion level of CD8 T-cells, *Q*, explicitly, following Johnson et al.[3]. $Q$ is dependent on the cumulative level of antigenic stimulation rather than the instantaneous level, in accordance with experiments[4]. The equations for model #3 are:

$$
\begin{aligned}
\frac{dT}{dt} &= \lambda - \beta' TI - d_T T \\
\frac{dI}{dt} &= (1 - f_D)\beta' TI - K^* E^* I - d_I I \\
\frac{dD}{dt} &= f_D \beta' TI - d_D D \\
\frac{dE^*}{dt} &= \lambda_E^* + \alpha_E E^* \frac{I}{\theta_E + I} - \xi E^* \frac{Q^n}{\phi^n + Q^n} - d_E E^* \\
\frac{dQ}{dt} &= \kappa \frac{I}{\theta_E + I} - d_Q Q \\
\frac{dK^*}{dt} &= \omega (1 - K^*)
\end{aligned} \tag{S2}
$$

Here, $\kappa$ is the maximum rate at which the exhaustion level rises, $d_Q$ is the rate constant for the reversal of exhaustion, $\xi$ is the maximum rate at which the effector cells suffer exhaustion and $\phi$ is the corresponding half-maximal saturation constant. The Hill coefficient for exhaustion, $n$, is 1[3].

Model #4 is identical to model #3, except for the Hill coefficient, which is set to 4[5].

In model #5, following Desikan et al.[5], we explicitly accounted for antigen-dependent enhanced recruitment of effector cells. This changes the equation for CD8 T-cells to $\frac{dE^*}{dt} = \lambda_E^* + (\rho_E + \alpha_E E^*)\frac{I}{\theta_E + I} - d_E E^*$, where $\rho_E$ is the maximum rate of antigen-dependent recruitment. For simplicity, we considered the same half-maximal saturation constant, $\theta_E$, for both antigen-dependent recruitment and proliferation events. The other equations in the model are identical to those in model #3.

Fits of the above models to the data are in Fig. S2 – S5, and the corresponding parameter estimates are in Table. S2 – S5.

#### TEXT S2. DERIVATION OF $\tau_{\max}$

In this section, we present the derivation of the time, $\tau_{\max}$, at which the antigen load in the CD4 T-cell culture in our *ex vivo* model setup (equation (11), main text) peaks.

Because virus-induced cytopathicity is negligible in these assays[6], we simplified equation (13) by setting $\delta \approx 0$, so that

$$\hat{V}(\tau) \approx \frac{\hat{V}_0}{2\alpha_n}\left((\rho+\alpha_n)e^{-\tau(\rho-\alpha_n)/2} - (\rho-\alpha_n)e^{-\tau(\rho+\alpha_n)/2}\right) \quad \text{(S3)}$$

where $\alpha_n = \sqrt{\rho^2 + 4(1-f)\hat{\beta}\hat{T}_0\phi}$. We substituted equation (S3) into the differential equation for $\hat{T}$, $\frac{d\hat{T}}{d\tau} = -\hat{\beta}\hat{T}\hat{V}$ (equation (13), main text), solving which yielded

$$\hat{T}(\tau) = \hat{T}_0 \exp\left(-\frac{4\hat{\beta}\rho\hat{V}_0}{\rho^2-\alpha^2}\right)\times \exp\left(\frac{\hat{V}_0\hat{\beta}}{\alpha_n}\left[\frac{\rho+\alpha_n}{\rho-\alpha_n}\exp\left(-\frac{\tau(\rho-\alpha_n)}{2}\right) - \frac{\rho-\alpha_n}{\rho+\alpha_n}\exp\left(-\frac{\tau(\rho+\alpha_n)}{2}\right)\right]\right) \quad \text{(S4)}$$

From Table S1, $\hat{\beta} = 10^{-8}$ mL copies$^{-1}$ d$^{-1}$, $\rho = 0.36$ d$^{-1}$, $\alpha_n \sim 1$ d$^{-1}$ and $\hat{V}_0 = 10^{2.86}$ copies mL$^{-1}$, which implied that $\exp\left(-\frac{4\hat{\beta}\rho\hat{V}_0}{\rho^2-\alpha^2}\right) \sim 1$. Further, $\exp(-\tau(\rho+\alpha_n)/2)$ vanishes quickly compared to $\exp(-\tau(\rho-\alpha_n)/2)$. The expression for $\hat{T}(\tau)$ in equation (S4) therefore simplified to

$$\hat{T}(\tau) \approx \hat{T}_0 \times \exp\left(\frac{\hat{V}_0\hat{\beta}}{\alpha_n}\frac{\rho+\alpha_n}{\rho-\alpha_n}\exp(-\tau(\rho-\alpha_n)/2)\right) \quad \text{(S5)}$$

From $\hat{T}(\tau)$, the effective reproductive ratio of the virus, $\mathcal{R}_{\text{eff}}$, followed as

$$\mathcal{R}_{\text{eff}} = \mathcal{R}_0 (1-\epsilon),\ \epsilon = 1 - \frac{\hat{T}(\tau)}{\hat{T}_0} \tag{S6}$$

where $\epsilon$ is the fraction of the target cells infected by time $\tau$, and $\mathcal{R}_0$ is the basic reproductive ratio, estimated at the start of the infection[7]. $\mathcal{R}_{\text{eff}}$ is the number of new infected cells that a single infected cell can give rise to in its lifetime[7, 8]. At $\tau = 0$, the infection spreads at the rate $\mathcal{R}_0$. As $\epsilon$ increases due to the depletion of target cells, the infection slows down. At the target cell concentration when $\mathcal{R}_{\text{eff}}$ drops below 1, the infection starts subsiding[7]. Thus, $\tau_{\max}$ is the time when $\mathcal{R}_{\text{eff}} = 1$. In other words, $\epsilon = 1 - 1/\mathcal{R}_0$ at $\tau = \tau_{\max}$.

To estimate $\mathcal{R}_0$, we used the next-generation matrix method. From our system of equations for the CD4 T-cell culture (equation (11), main text), we recognized that the infection subsystem (see [8]) will have the following equations for $\hat{S}$ and $\hat{V}$:

$$\begin{aligned} \frac{d\hat{S}}{d\tau} &= (1-f)\hat{\beta}\hat{T}\hat{V} - \rho\hat{S} \\ \frac{d\hat{V}}{d\tau} &= \phi\hat{S} - \delta\hat{V} \end{aligned} \tag{S7}$$

$F$ and $W$ denote the transmission and transition matrices, respectively, at the infection-free steady state [8], and are defined as

$$F = \left.\frac{\partial \mathcal{F}_i}{\partial x_j}\right|_0 \text{ and } W = \left.\frac{\partial \mathcal{W}_i}{\partial x_j}\right|_0 \tag{S8}$$

where $\vec{x} = \left(\hat{S}, \hat{V}\right)$. $\mathcal{F}_i$ denotes the new infection events in the $i^{\text{th}}$ compartment. $\mathcal{W}_i = \mathcal{W}_i^- - \mathcal{W}_i^+$, with $\mathcal{W}_i^+$ and $\mathcal{W}_i^-$ the transition events into and out of the $i^{\text{th}}$ compartment, respectively. The subscript 0 in equation (S8) denotes the evaluation of the partial derivatives at the infection-free steady state ($\hat{T} = \hat{T}_0$ and $\hat{S} = \hat{V} = 0$). This yields $\mathcal{F}_1 = (1-f)\hat{\beta}\hat{T}\hat{V}$, $\mathcal{F}_2 = 0$, $\mathcal{W}_1 = \rho\hat{S}$ and $\mathcal{W}_2 = \delta\hat{V} - \phi\hat{S}$ respectively, and hence

$$F = \begin{pmatrix} 0 & (1-f)\hat{\beta}\hat{T}_0 \\ 0 & 0 \end{pmatrix}, \; W = \begin{pmatrix} \rho & 0 \\ -\phi & \delta \end{pmatrix} \tag{S9}$$

$\mathcal{R}_0$ is the spectral radius of the matrix $FW^{-1}$:

$$FW^{-1} = \begin{pmatrix} 0 & (1-f)\hat{\beta}\hat{T}_0 \\ 0 & 0 \end{pmatrix} \times \frac{1}{\rho\delta}\begin{pmatrix} \delta & 0 \\ \phi & \rho \end{pmatrix} = \begin{pmatrix} \dfrac{(1-f)\hat{\beta}\phi}{\rho\delta}\hat{T}_0 & \dfrac{(1-f)\hat{\beta}}{\delta}\hat{T}_0 \\ 0 & 0 \end{pmatrix} \tag{S10}$$

Thus,

$$\mathcal{R}_0 = (1-f)\hat{\beta}\phi\hat{T}_0/\rho\delta \tag{S11}$$

Combining equations (S5), (S6) and (S11) then yielded

$$\tau_{\max} = \frac{2}{\alpha_n - \rho}\ln\left[\frac{\alpha_n}{\hat{V}_0\hat{\beta}}\frac{\rho+\alpha_n}{\rho-\alpha_n}\ln\left(\frac{\rho\delta}{(1-f)\hat{\beta}\phi\hat{T}_0}\right)\right] \tag{S12}$$

Estimates of $\mathcal{R}_0$, $\epsilon$ and $\tau_{\max}$ for the parameter values employed are presented in Table S7.

### SUPPLEMENTARY FIGURES

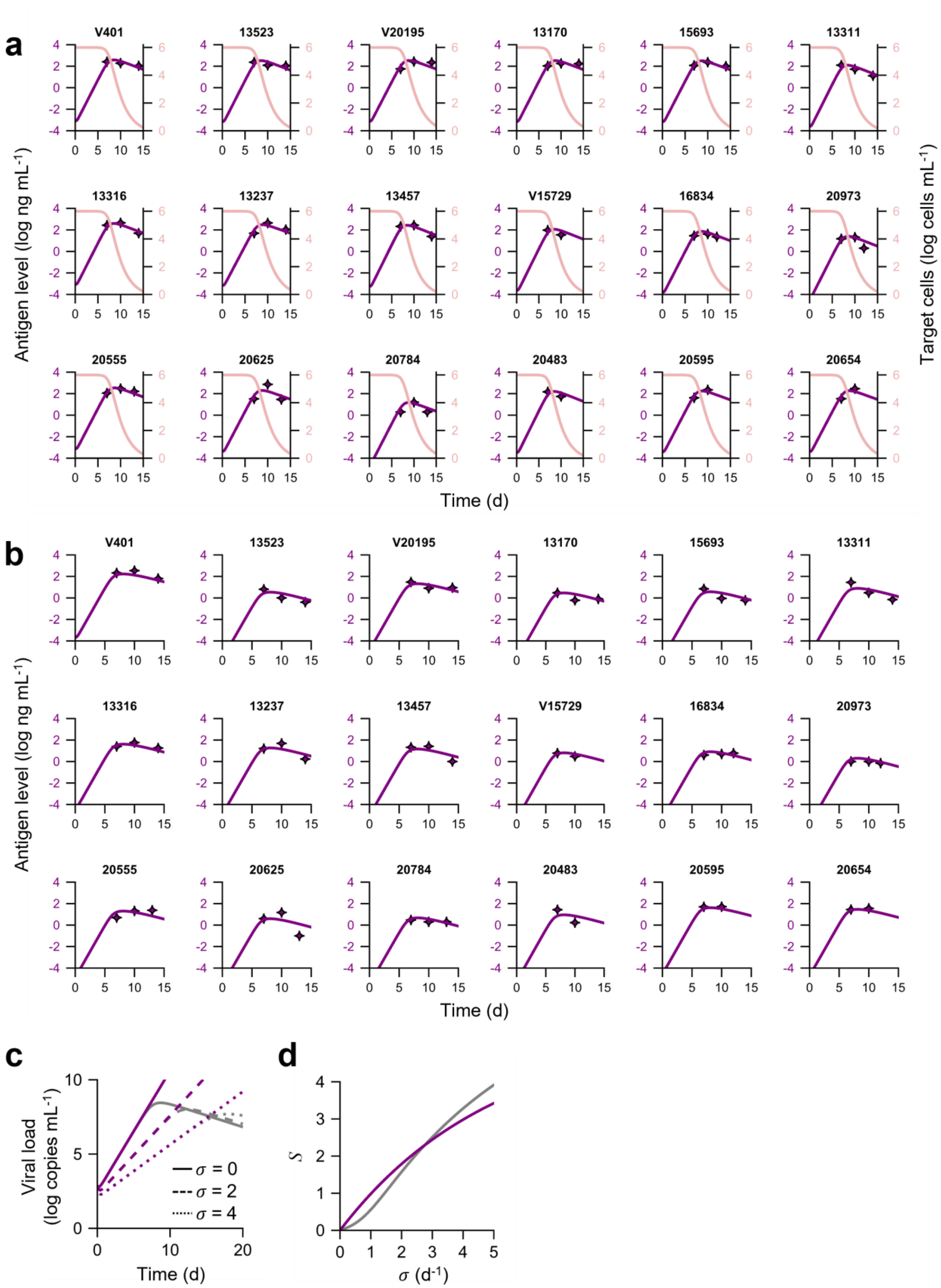

**Fig. S1: The *ex vivo* model predictions and fits. (a)** Fits (lines) of the *ex vivo* model (equation (11), main text) to antigen load data (symbols) from CD4 T-cell cultures of 18 samples. Sample IDs are presented on the top of the corresponding panels. Antigen p27 level is assumed to be $\mu\hat{V}$, where $\hat{V}$ is the viral load and $\mu$ is the amount of antigen per copy of virion. $\mu$ and $\rho$ were identifiable and were estimated to be $6.2\times10^{-7}$ ng copies$^{-1}$ and 0.36 d$^{-1}$, respectively. The pink curves plot the corresponding target cell concentrations. **(b)** Fits of the *ex vivo* model to the 1:1 CD4 and CD8 T-cell co-cultures of 18 samples. Sample IDs are presented on the top of corresponding panels. Estimated $\rho$ from fits to CD4 T-cell cultures were used and $\sigma$ was adjusted to fit the model. **(c)** Estimates of viral load in the cultures by equation (13) from main text (purple) and numerical integration of system in equation (11) from main text (gray). The CD4 T-cell culture corresponds to $\sigma = 0$, while the other cases are co-cultures. **(d)** Estimates of the suppressive capacity calculated from the equation (14) from main text (purple) and the numerical integration of system (equation (11), main text) (gray).

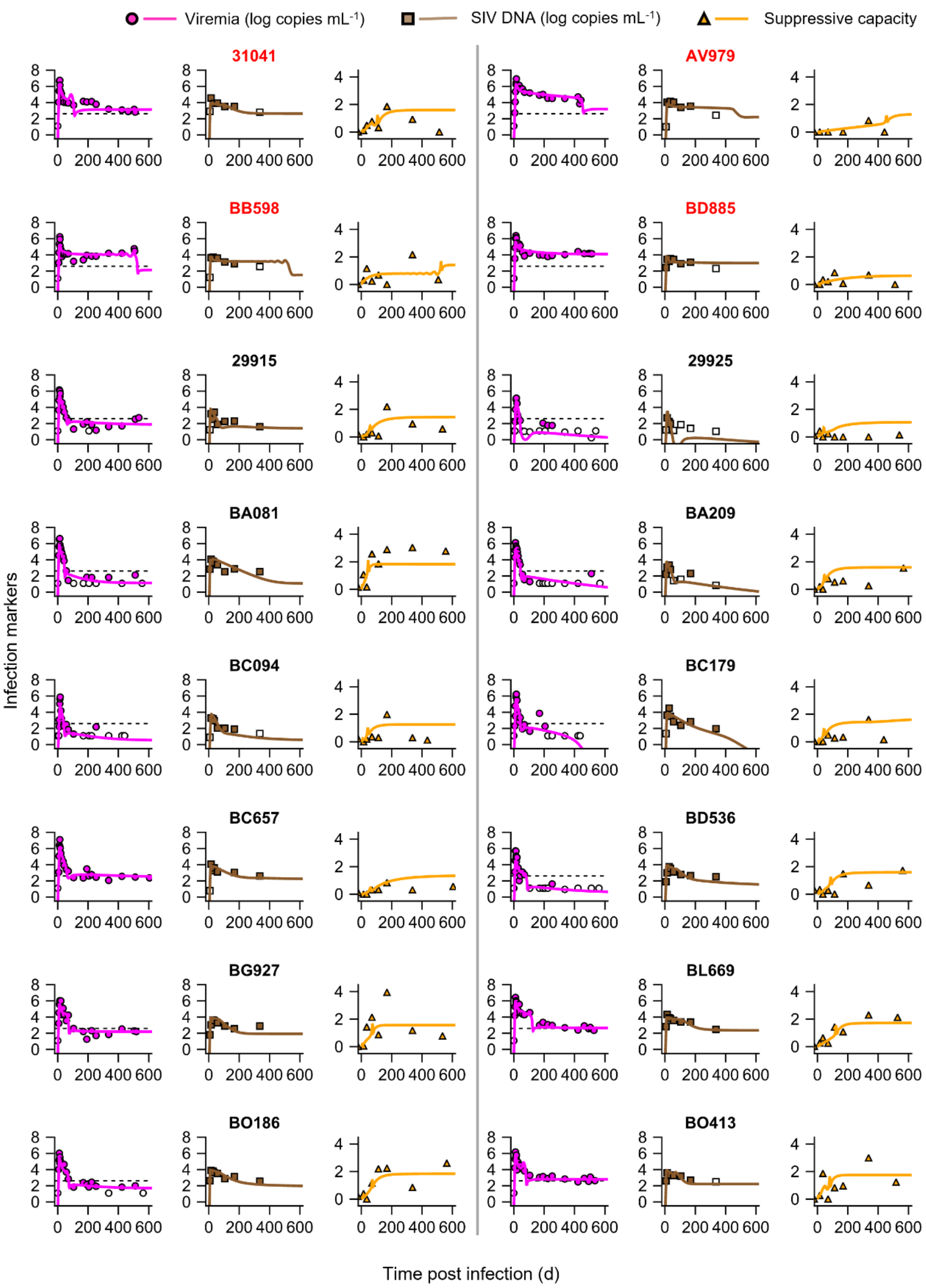


**Fig. S2: Fits of model #2 to data.** Model predictions (lines) from simultaneous fitting of model #2 (Methods; Table S1) to all the three datasets (symbols), namely, viremia (magenta), SIV DNA (brown) and suppressive capacity (yellow). Macaques highlighted in red are progressors while the rest are controllers. Empty symbols are observations below the limit of detection. The parameter estimates resulting in these fits are in Table S2.

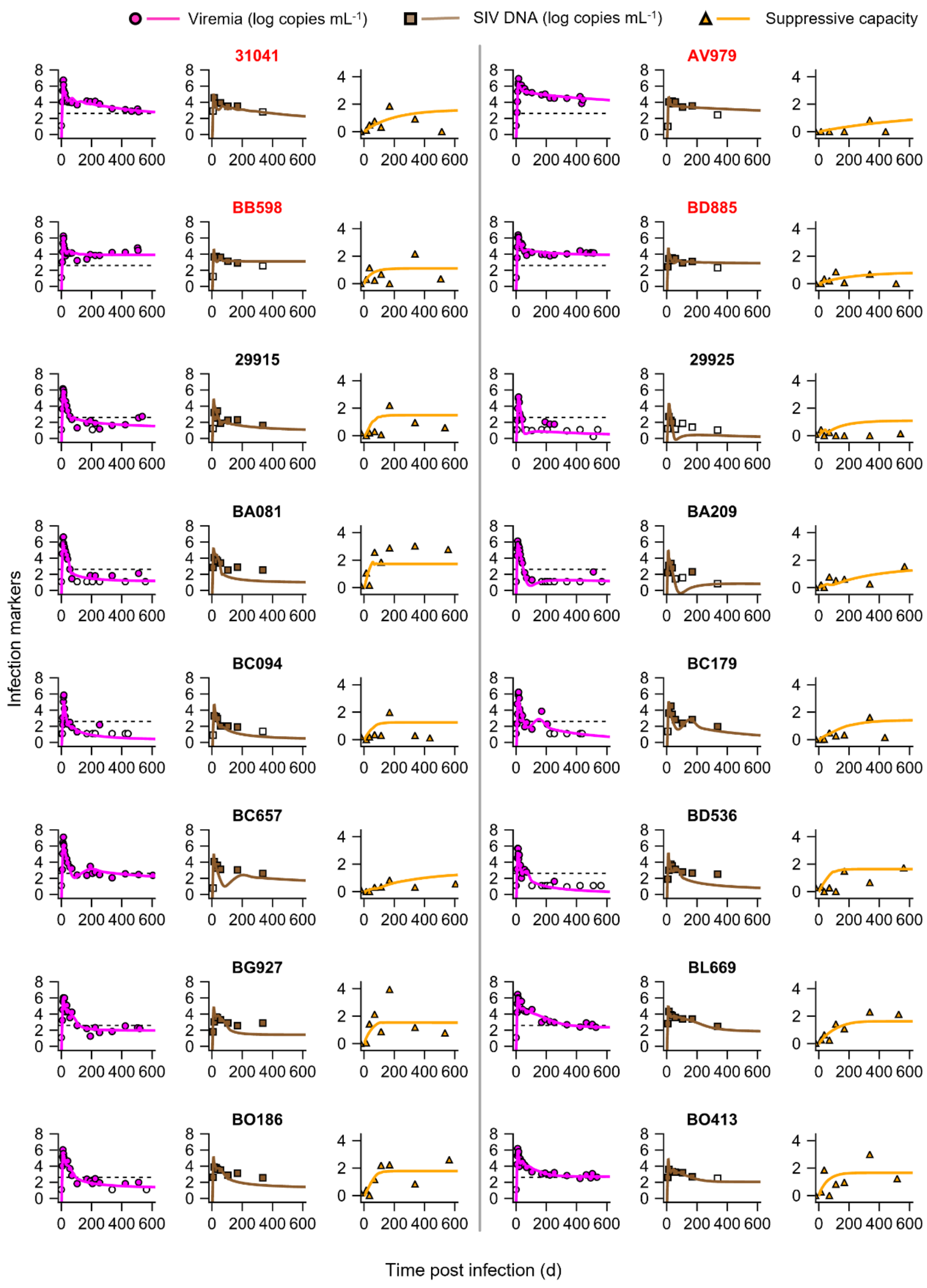


**Fig. S3: Fits of model #3 to data.** Model predictions (lines) from simultaneous fitting of model #3 (Methods; Table S1) to all the three datasets (symbols), namely, viremia (magenta), SIV DNA (brown) and suppressive capacity (yellow). Macaques highlighted in red are progressors while the rest are controllers. Empty symbols are observations below the limit of detection. The parameter estimates resulting in these fits are in Table S3.

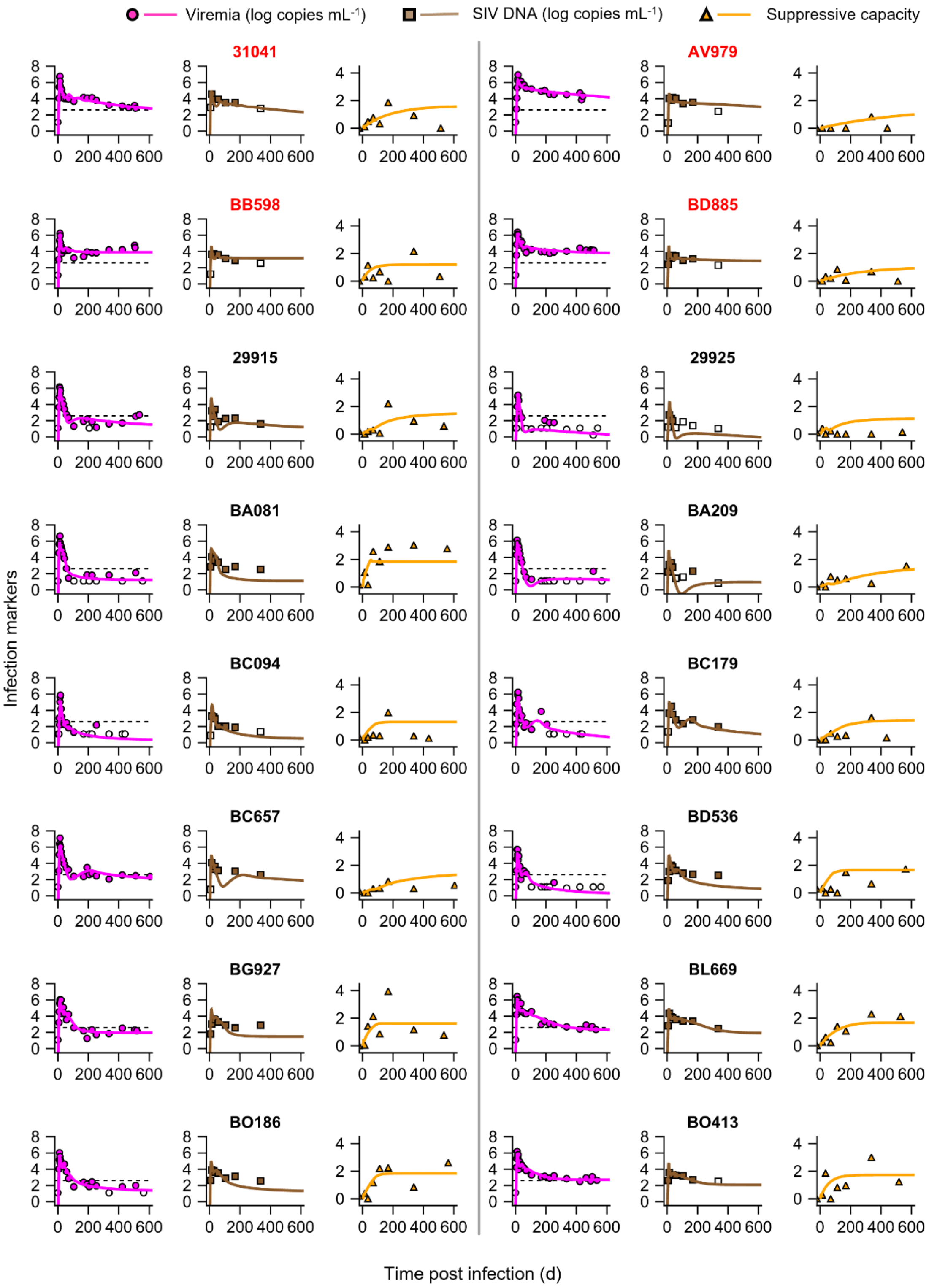


**Fig. S4: Fits of model #4 to data.** Model predictions (lines) from simultaneous fitting of model #4 (Methods; Table S1) to all the three datasets (symbols), namely, viremia (magenta), SIV DNA (brown) and suppressive capacity (yellow). Macaques highlighted in red are progressors while the rest are controllers. Empty symbols are observations below the limit of detection. The parameter estimates resulting in these fits are in Table S4.

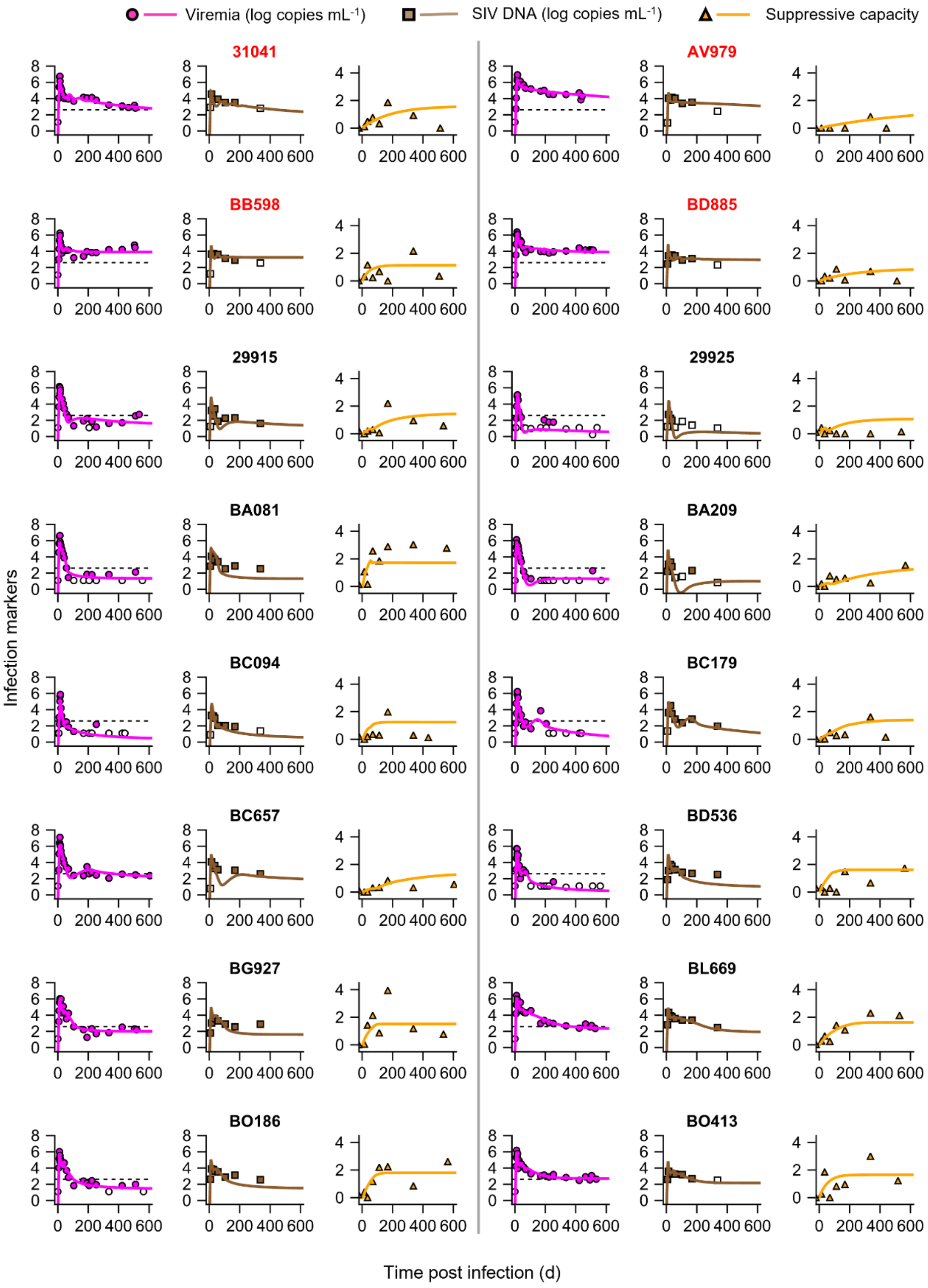


**Fig. S5: Fits of model #5 to data.** Model predictions (lines) from simultaneous fitting of model #5 (Methods; Table S1) to all the three datasets (symbols), namely, viremia (magenta), SIV DNA (brown) and suppressive capacity (yellow). Macaques highlighted in red are progressors while the rest are controllers. Empty symbols are observations below the limit of detection. The parameter estimates resulting in these fits are in Table S5.

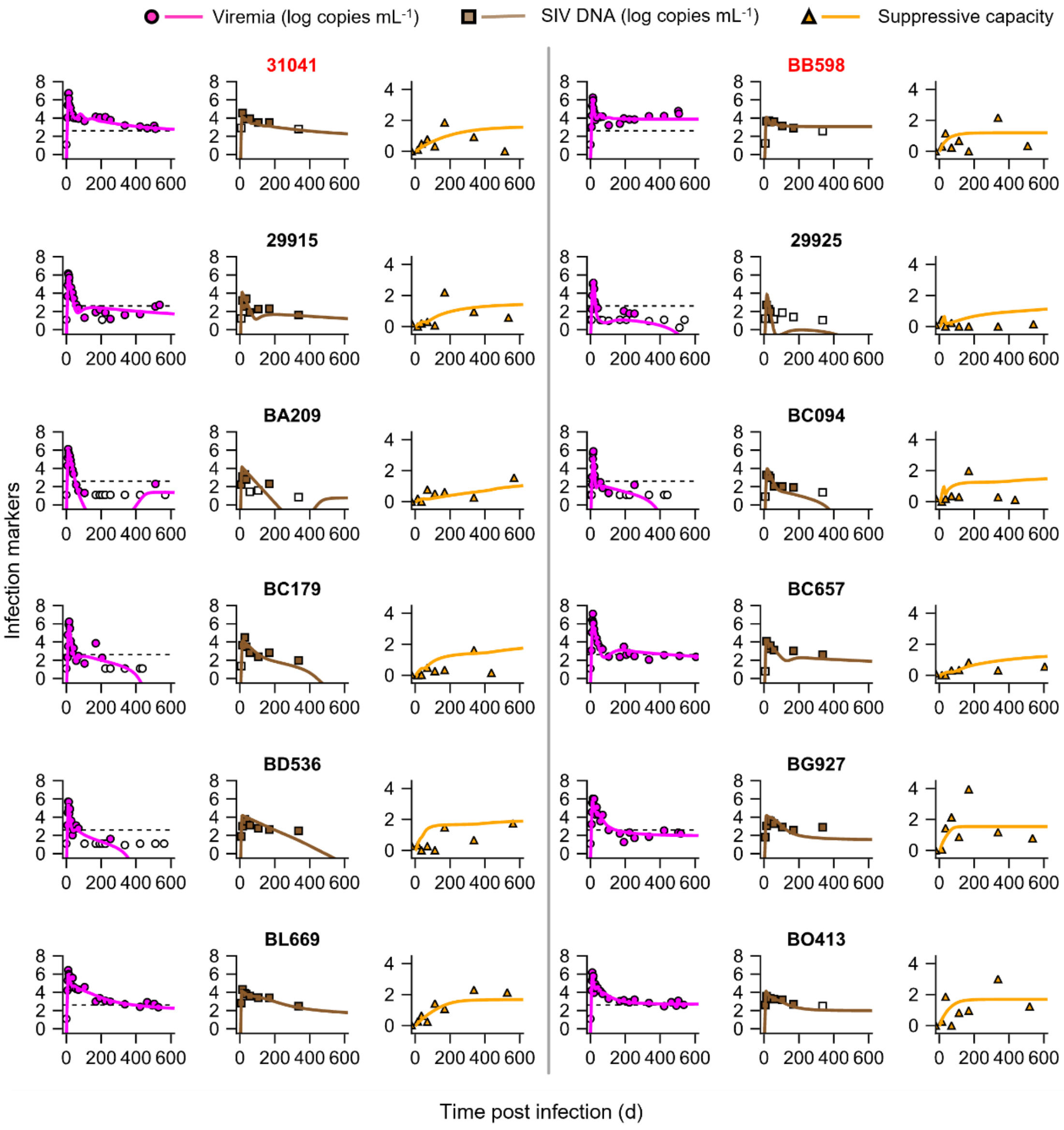


**Fig. S6: Fits of the best-fit model to data.** Model predictions (lines) from simultaneous fitting of the best-fit model (Methods; Table S1) to all the three datasets (symbols), namely, viremia (magenta), SIV DNA (brown) and suppressive capacity (yellow), shown for 12 of 16 macaques. Plots for the remaining 4 macaques are presented in Fig. 2. Macaques highlighted in red are progressors while the rest are controllers. Empty symbols are observations below the limit of detection. The parameter estimates resulting in these fits are detailed in Table 1 of the main text and Table S6.

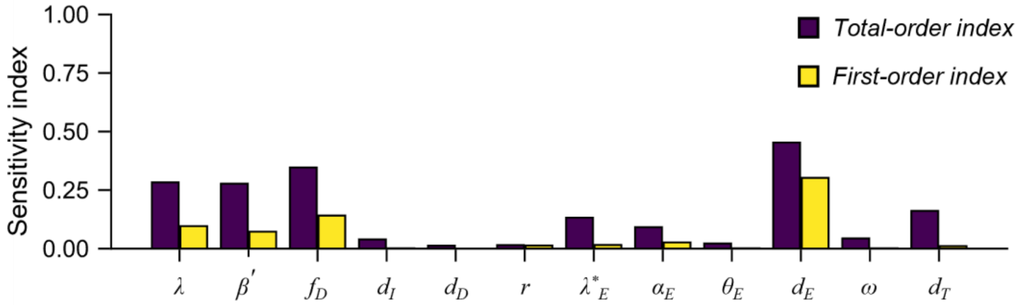


**Fig. S7: Sensitivity analysis.** Sensitivity of the set-point viral load predicted by the best-fit model to its parameters estimated using Sobol's method.

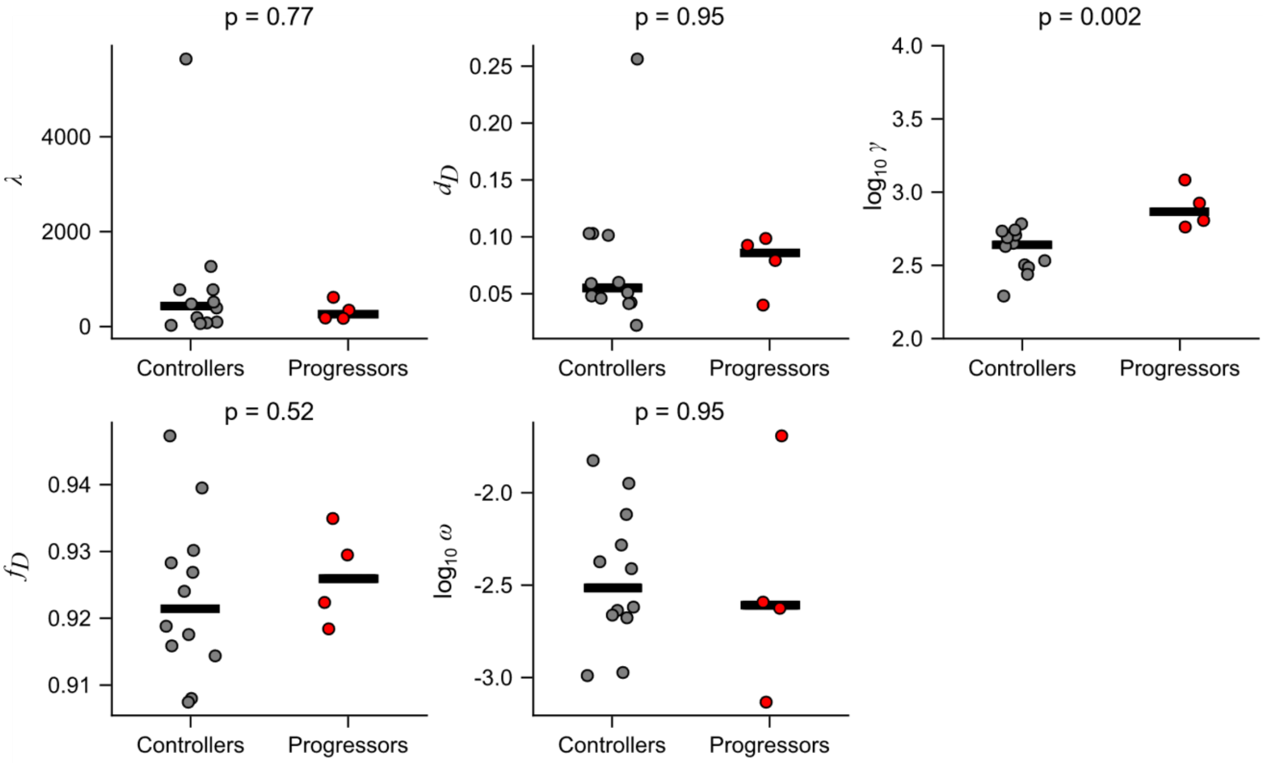


**Fig. S8: Comparison of parameters estimated by model #1.** Parameters estimated for all the individuals are grouped based on their control status – controllers vs. progressors – and compared. Presented here are five parameters ($\lambda$, $d_D$, $\alpha_E$, $f_D$ and $\log_{10}\omega$). The others are in Fig. 3. Mann-Whitney U test was used to estimate the significance levels.

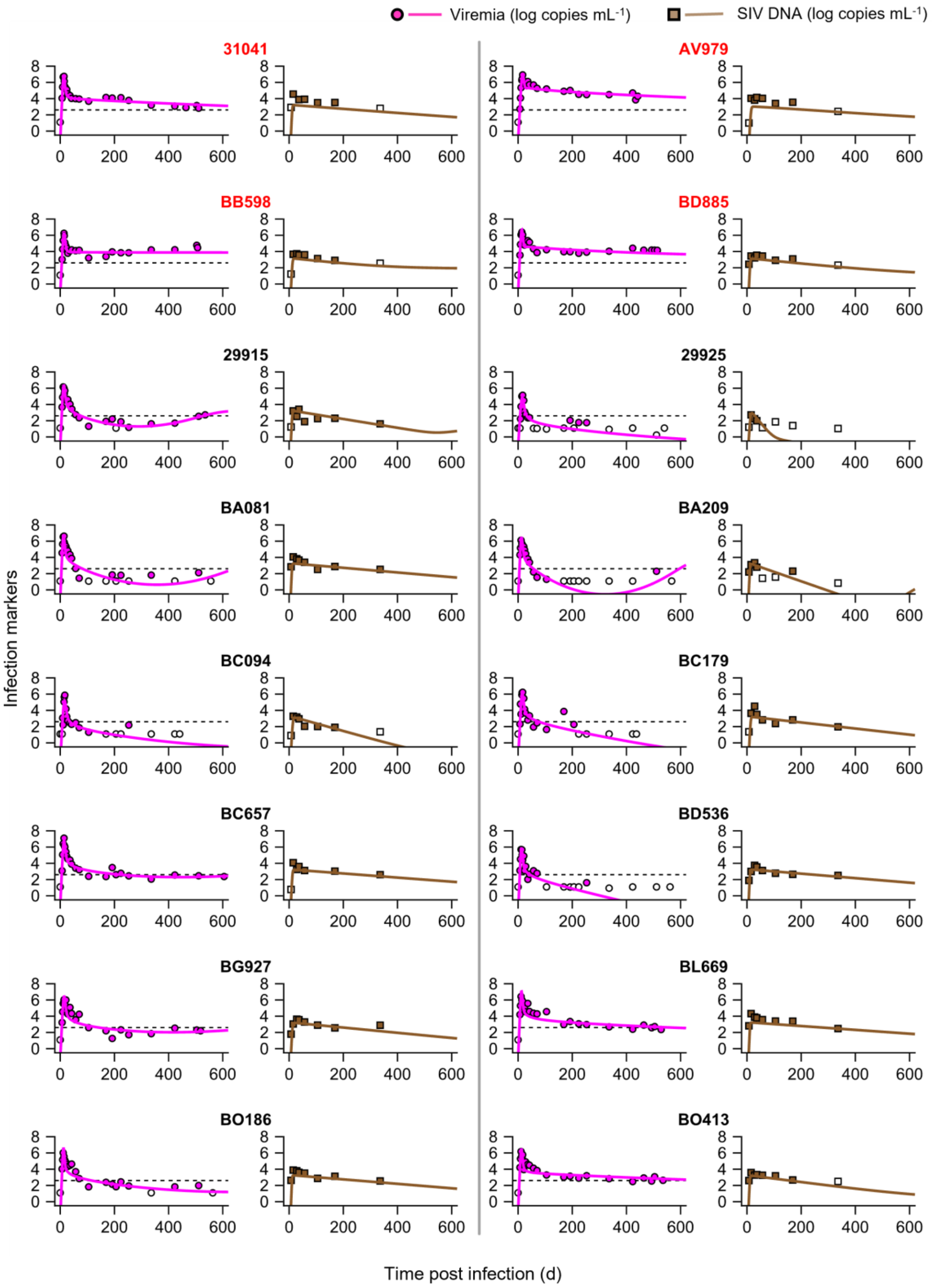


**Fig. S9: Fits of the model that does not incorporate suppressive capacity measurements to data.** Model predictions (lines) from simultaneous fitting of model #6 (Methods; Table S1) to all the two virological datasets (symbols), namely, viremia (magenta) and SIV DNA (brown). Macaques highlighted in red are progressors while the rest are controllers. Empty symbols are observations below the limit of detection. The parameter estimates resulting in these fits are in Table S8.

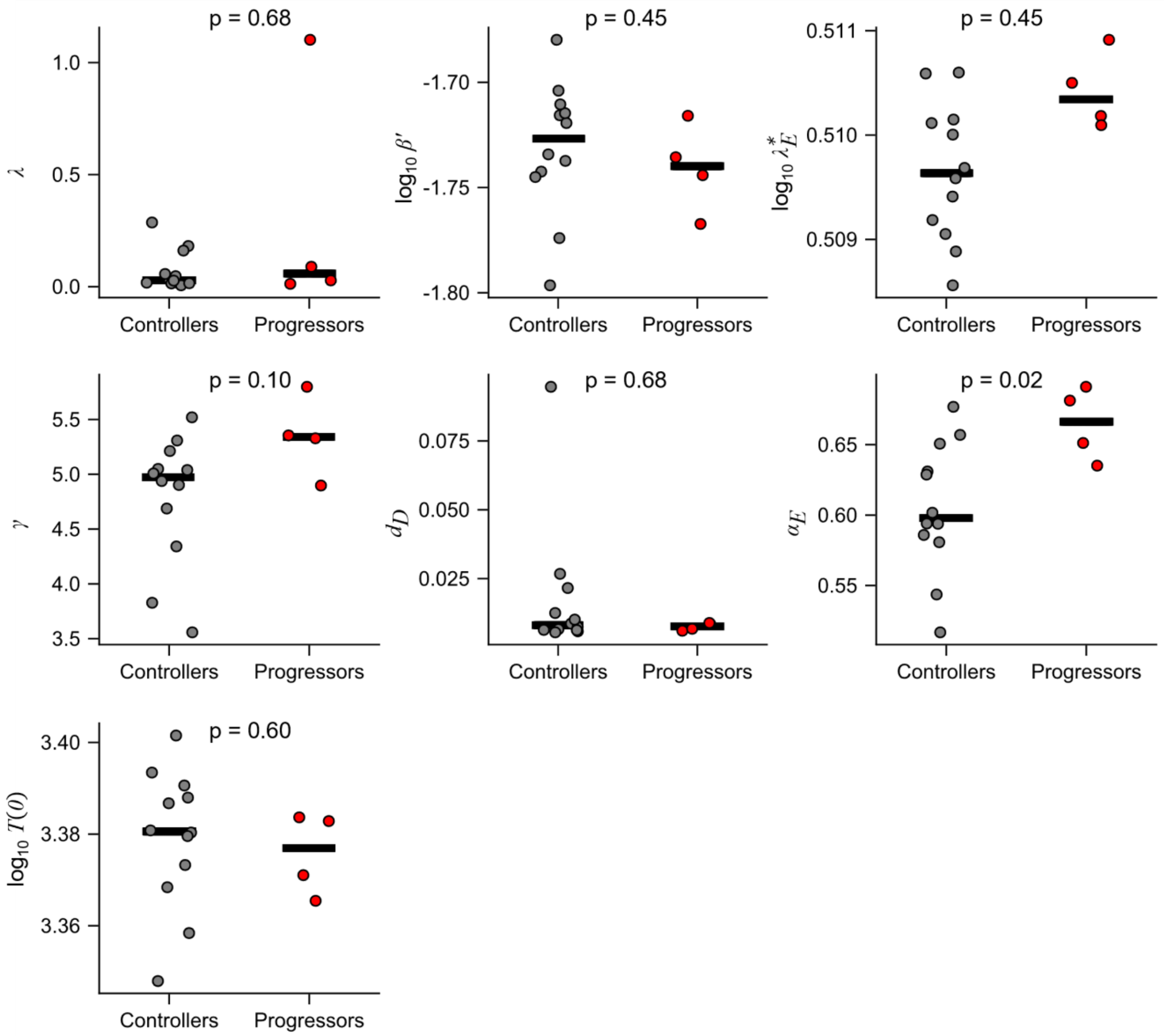


**Fig. S10: Comparison of parameters estimated by the model that does not incorporate suppressive capacity measurements for fitting.** Parameters estimated for all the individuals are grouped based on their control status – controllers vs. progressors – and compared. Mann-Whitney U test was used to estimate the significance levels.

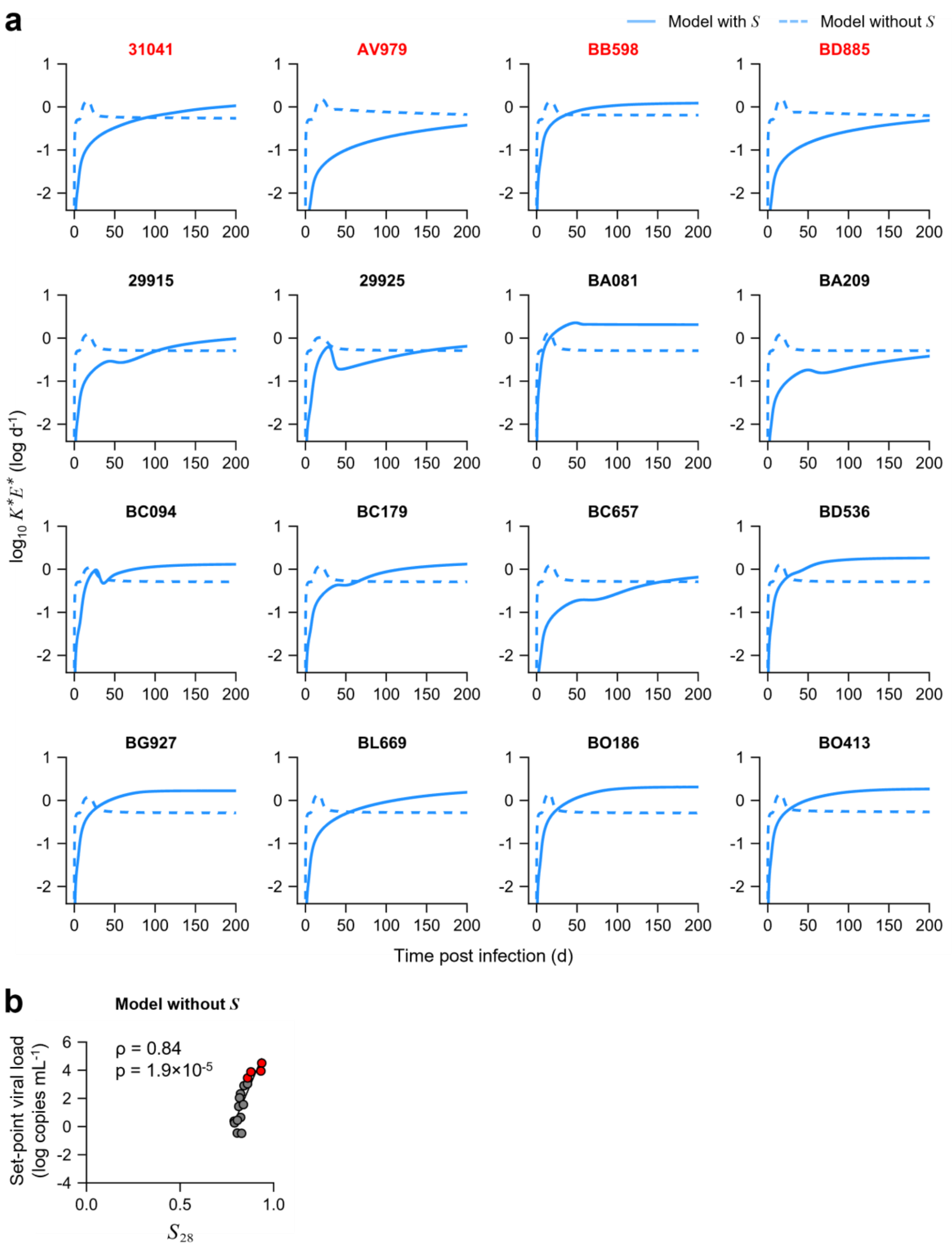


**Fig. S11: Comparison of CD8 T-cell killing rate between model with and without suppressive capacity.** **(a)** Effector response dynamics of CD8 T-cells, given by $K^*E^*$, predicted for the macaques by the best-fit model (solid) and model #6, which does not incorporate suppressive capacity measurements for fitting (dashed). **(b)** Correlation plot between $S_{28}$ and set-point viral load as predicted by model #6. Gray symbols are controllers, while red symbols are progressors. Spearman's ρ was calculated for assessing the correlation. Note that here the set-point viral load increases with $S_{28}$, which is the opposite of what is expected.

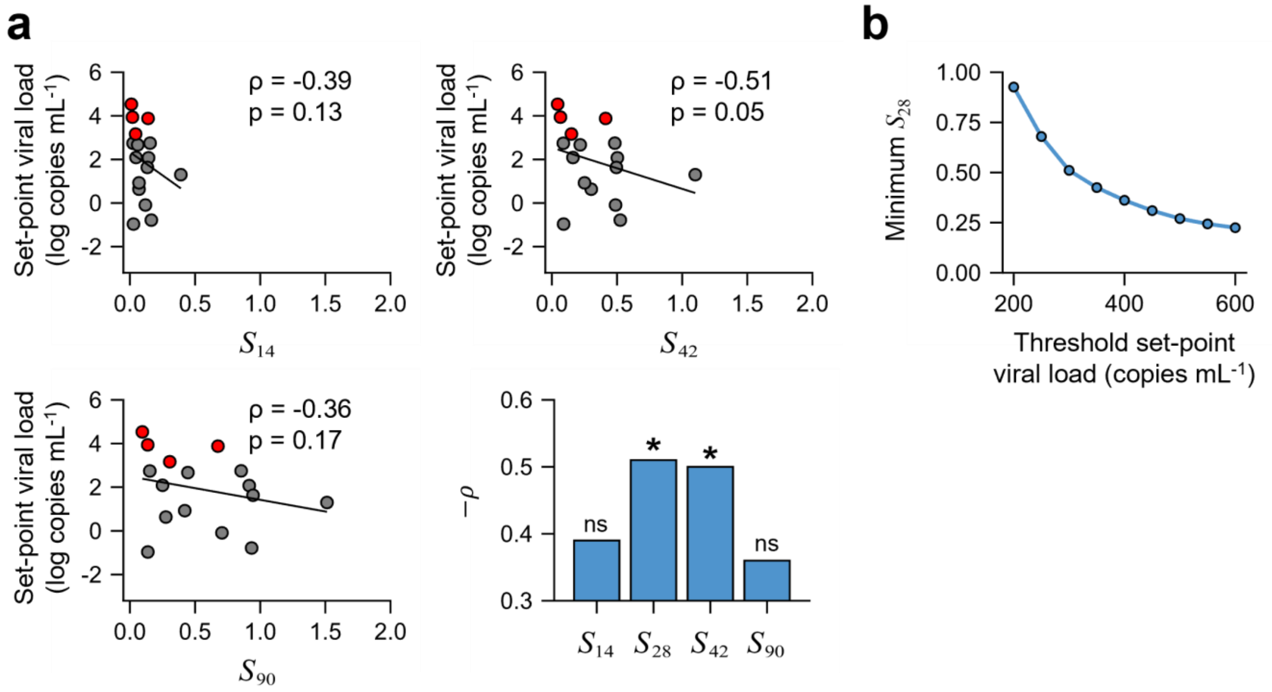


**Fig. S12: Robustness of correlate. (a) Sensitivity to duration for evaluating the early CD8 T-cell responses.** Correlation between set-point viral load and AUC of suppressive capacity averaged over 14, 42 and 90 days post infection, respectively, for the 16 macaques. Gray symbols are controllers, while red symbols are progressors. The bar plot at the bottom right presents the predicted correlation between set-point viral load and the time-averaged area-under-the-curve of $S$ estimated for different durations. Asterisks represent significant correlations with $p<0.05$; ns: not significant. **(b) Minimum $S_{28}$ required for control increases with a stricter definition of control.** The minimum $S_{28}$ estimated to be required for 95% likelihood of control as a function of the threshold viral load for control. Spearman's ρ was calculated for assessing the correlations.

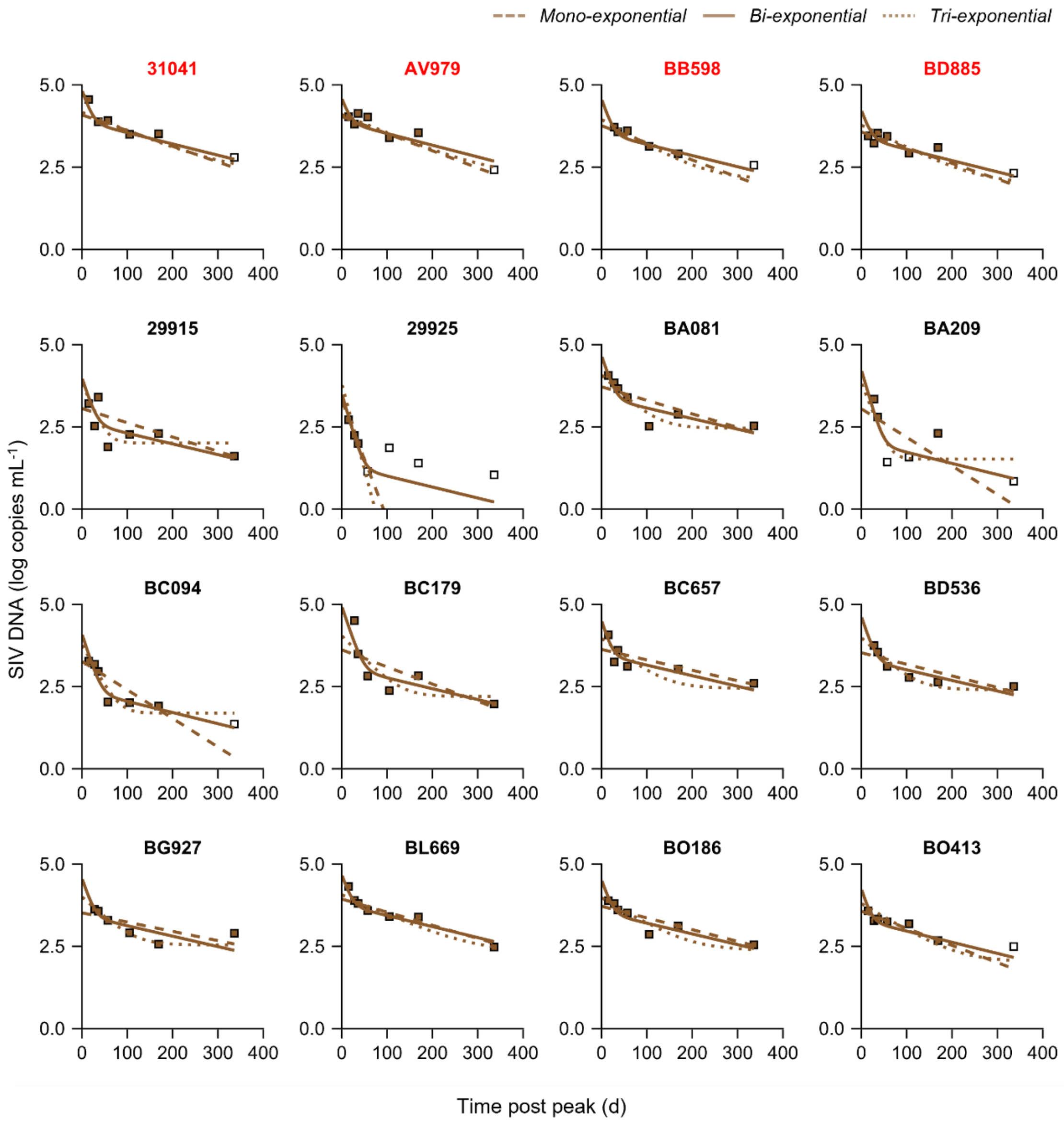


**Fig. S13: Identifying number of phases of SIV DNA.** Mono- (dashed), bi- (solid), and tri-exponential (dotted) curves are fitted to longitudinal SIV DNA data post the peak in the measurements. Empty symbols are below detection limit. Data were fit in Monolix (Methods; main text). The bi-exponential curve explained the data best (BICs: 164.48 for the mono-exponential curve; 140.36 for the bi-exponential curve; and 171.13 for the tri-exponential curve).

### SUPPLEMENTARY TABLES

| Model | Description | BIC | Observation | Figure | Parameters |
|---|---|---|---|---|---|
| 1 | Best-fit model (equations (1) – (10) in main text) | 1505 | Explains the data best | Fig. 2; Fig. S6 | Table 1; Table S6 |
| 2 | Model with exhaustion dependent on instantaneous antigen level (Equation (S1)) | 1546 | Model fits do not capture the data any better than the main model | Fig. S2 | Table S2 |
| 3 | Model with exhaustion dependent on cumulative antigen level; Hill coefficient $n$ = 1 (Equation (S2)) | 1563 | Model fits do not capture the data any better than the main model | Fig. S3 | Table S3 |
| 4 | Model with exhaustion dependent on cumulative antigen level; Hill coefficient $n$ = 4 (Equation (S2)) | 1557 | Model fits do not capture the data any better than the main model | Fig. S4 | Table S4 |
| 5 | Main model with added antigen-dependent effector CD8 T-cell recruitment | 1544 | Model fits do not capture the data any better than the main model | Fig. S5 | Table S5 |
| 6 | Main model fitted without suppressive capacity dataset and constant $k$ | 1168* | Effector CD8 T-cell dynamics are captured worse than the main model | Fig. S9 | Table S8 |

**Table S1: Comparison of different models fitted to the data.** Every model fit to the data is summarized, comparing the BICs of fits. *This fit does not include suppressive capacity datasets and hence cannot be compared with other models.

| Parameter (Units) | Fixed effect | Random effect |
|---|---|---|
| $\lambda$ (cells mL$^{-1}$ d$^{-1}$) | 210 (56.6) | 0.76 (0.22) |
| $\log_{10}\beta'$ (log mL cells$^{-1}$ d$^{-1}$) | -2.39 (0.05) | 0.06 (0.04) |
| $f_D$ | 0.95 | - |
| $\log_{10}\lambda_E{}^*$ (log d$^{-2}$) | -0.03 (0.07) | 0.22 (0.06) |
| $d_I$ (d$^{-1}$) | 0.10 | - |
| $d_D$ (d$^{-1}$) | 0.06 (0.02) | 0.91 (0.22) |
| $\gamma$ (cells$^{-1}$) | 517 (169) | 1.13 (0.27) |
| $\alpha_E$ (d$^{-1}$) | 1.00 | - |
| $\theta_E$ (cells mL$^{-1}$) | 0.10 | - |
| $\alpha_R$ (d$^{-1}$) | 0.02 (0.04) | 0.94 (2.83) |
| $\theta_X$ (cells mL$^{-1}$) | 5.00 | - |
| $d_E$ (d$^{-1}$) | 1.00 | - |
| $\log_{10}\omega$ (log d$^{-1}$) | -2.15 (0.13) | 0.37 (0.09) |
| $\log_{10}T(0)$ (log cells mL$^{-1}$) | 3.94 (0.05) | 0.01 (0.01) |

**Table S2: Population parameter estimates for model #2.** The fixed and random effects of each parameter is provided along with respective standard errors in parentheses. In addition to the parameters fixed in model #1, $f_D$ is fixed to 0.95 and $\theta_X$ is fixed to 5 d$^{-1}$ (ref [1, 9]).

| **Parameter (Units)** | **Fixed effect** | **Random effect** |
|---|---|---|
| $\lambda$ (cells $mL^{-1}$ $d^{-1}$) | $8.91\times10^{3}$ ($3.42\times10^{3}$) | 1.13 (0.24) |
| $\log_{10}\beta'$ (log mL $cells^{-1}$ $d^{-1}$) | -4.10 (0.13) | 0.02 (0.12) |
| $f_D$ | 0.95 | - |
| $\log_{10}\lambda_E{}^*$ (log $d^{-2}$) | -0.189 (0.525) | 0.27 (0.29) |
| $d_I$ ($d^{-1}$) | 0.10 | - |
| $d_D$ ($d^{-1}$) | 1.72 (1.35) | 0.75 (0.58) |
| $\gamma$ ($cells^{-1}$) | 21.3 (13.2) | 0.78 (0.28) |
| $\alpha_E$ ($d^{-1}$) | 1.66 (0.35) | 0.02 (0.17) |
| $\theta_E$ (cells $mL^{-1}$) | 0.10 | - |
| $\xi$ ($d^{-1}$) | 1.83 (4.08) | 0.17 (0.21) |
| $\phi$ ($mL^{-1}$) | 2.00 | - |
| $d_E$ ($d^{-1}$) | 1.00 | - |
| $\kappa$ ($d^{-1}$) | 1.00 | - |
| $d_Q$ ($d^{-1}$) | 0.52 (1.45) | 0.19 (0.82) |
| $\log_{10}\omega$ (log $d^{-1}$) | -2.48 (0.36) | 0.45 (0.14) |
| $\log_{10}T(0)$ (log cells $mL^{-1}$) | 5.64 (0.13) | 0.01 (0.01) |

**Table S3: Population parameter estimates for model #3.** The Hill coefficient for the exhaustion rate, $n$=1. The fixed and random effects of each parameter is provided along with respective standard errors in parentheses. In addition to the parameters fixed in model #1, $f_D$ is fixed to 0.95, $\phi$ is fixed to 2 $mL^{-1}$ and $\kappa$ is fixed to 1 $d^{-1}$ (ref [1, 3, 9]).

| **Parameter (Units)** | **Fixed effect** | **Random effect** |
|---|---|---|
| $\lambda$ (cells $mL^{-1}$ $d^{-1}$) | $3.41\times10^{3}$ ($4.60\times10^{3}$) | 1.17 (0.30) |
| $\log_{10}\beta'$ (log mL $cells^{-1}$ $d^{-1}$) | -3.66 (0.52) | 0.03 (0.09) |
| $f_D$ | 0.95 | - |
| $\log_{10}\lambda_E{}^*$ (log $d^{-2}$) | 0.29 (0.29) | 0.16 (0.17) |
| $d_I$ ($d^{-1}$) | 0.10 | - |
| $d_D$ ($d^{-1}$) | 0.52 (0.92) | 0.76 (0.86) |
| $\gamma$ ($cells^{-1}$) | 63.6 (81.8) | 0.74 (0.36) |
| $\alpha_E$ ($d^{-1}$) | 1.23 (1.1) | 0.19 (0.40) |
| $\theta_E$ (cells $mL^{-1}$) | 0.10 | - |
| $\xi$ ($d^{-1}$) | 0.99 (1.72) | 0.28 (0.58) |
| $\phi$ ($mL^{-1}$) | 2.00 | - |
| $d_E$ ($d^{-1}$) | 1.00 | - |
| $\kappa$ ($d^{-1}$) | 1.00 | - |
| $d_Q$ ($d^{-1}$) | 0.01 (0.04) | 1.08 (5.81) |
| $\log_{10}\omega$ (log $d^{-1}$) | -2.37 (0.28) | 0.40 (0.11) |
| $\log_{10}T(0)$ (log cells $mL^{-1}$) | 5.22 (0.55) | 0.01 (0.01) |

**Table S4: Population parameter estimates for model #4.** The Hill coefficient for the exhaustion rate, $n$=4. The fixed and random effects of each parameter is provided along with respective standard errors in parentheses. In addition to the parameters fixed in model #1, $f_D$ is fixed to 0.95, $\phi$ is fixed to 2 $mL^{-1}$ and $\kappa$ is fixed to 1 $d^{-1}$ (ref [1, 3, 9]).

| Parameter (Units) | Fixed effect | Random effect |
|---|---|---|
| $\lambda$ (cells mL$^{-1}$ d$^{-1}$) | $3.13\times10^{3}$ ($2.27\times10^{3}$) | 1.20 (0.27) |
| $\log_{10}\beta'$ (log mL cells$^{-1}$ d$^{-1}$) | -3.65 (0.12) | 0.02 (0.06) |
| $f_D$ | 0.95 | - |
| $\log_{10}\lambda_E{}^*$ (log d$^{-2}$) | -0.23 (1.65) | 0.29 (1.01) |
| $d_I$ (d$^{-1}$) | 0.10 | - |
| $d_D$ (d$^{-1}$) | 0.45 (0.23) | 0.62 (0.53) |
| $\gamma$ (cells$^{-1}$) | 65.90 (30.60) | 0.77 (0.30) |
| $\rho_E$ (cells mL$^{-1}$ d$^{-2}$) | 0.25 (4.16) | 1.07 (6.74) |
| $\alpha_E$ (d$^{-1}$) | 0.66 (0.81) | 0.21 (0.52) |
| $\theta_E$ (cells mL$^{-1}$) | 0.10 | - |
| $d_E$ (d$^{-1}$) | 1.00 | - |
| $\log_{10}\omega$ (log d$^{-1}$) | -2.44 (0.20) | 0.42 (0.13) |
| $\log_{10}T(0)$ (log cells mL$^{-1}$) | 5.19 (0.11) | 0.01 (0.01) |

**Table S5: Population parameter estimates for model #5.** The fixed and random effects of each parameter is provided along with respective standard errors in parentheses. In addition to the parameters fixed in model #1, $f_D$ is fixed to 0.95 (ref [9]).

| **Macaque ID** | $\lambda$ | $f_D$ | $\log_{10}\lambda_E{}^*$ | $d_D$ | $r$ | $\alpha_E$ | $\log_{10}\omega$ |
|---|---|---|---|---|---|---|---|
| 29915 | 95.33 | 0.93 | 0.19 | 0.10 | 340.52 | 0.58 | -2.68 |
| 29925 | 77.18 | 0.95 | 0.18 | 0.26 | 318.82 | 0.86 | -2.64 |
| 31041 | 179.47 | 0.92 | 0.07 | 0.04 | 841.63 | 0.58 | -2.59 |
| AV979 | 344.91 | 0.93 | 0.06 | 0.08 | 1211.71 | 0.59 | -3.13 |
| BA081 | 5638.31 | 0.91 | 0.26 | 0.04 | 447.82 | 0.61 | -1.83 |
| BA209 | 25.91 | 0.92 | 0.31 | 0.05 | 508.71 | 0.50 | -2.99 |
| BB598 | 615.66 | 0.93 | -0.25 | 0.09 | 640.96 | 0.56 | -1.69 |
| BC094 | 391.44 | 0.94 | 0.23 | 0.10 | 306.32 | 0.83 | -2.37 |
| BC179 | 190.44 | 0.93 | 0.42 | 0.04 | 426.20 | 0.54 | -2.66 |
| BC657 | 62.97 | 0.93 | 0.20 | 0.05 | 607.85 | 0.59 | -2.97 |
| BD536 | 515.97 | 0.92 | 0.35 | 0.02 | 195.26 | 0.60 | -2.28 |
| BD885 | 172.81 | 0.92 | -0.23 | 0.10 | 577.93 | 0.55 | -2.63 |
| BG927 | 1266.57 | 0.92 | 0.04 | 0.06 | 488.93 | 0.69 | -2.12 |
| BL669 | 474.54 | 0.92 | 0.14 | 0.06 | 552.07 | 0.68 | -2.62 |
| BO186 | 775.64 | 0.91 | 0.27 | 0.05 | 274.69 | 0.72 | -2.41 |
| BO413 | 776.12 | 0.91 | -0.04 | 0.10 | 542.03 | 0.62 | -1.95 |

**Table S6: Individual parameter estimates for the best-fit model.** Fixed parameters are $d_I$, $\theta_E$, and $d_E$ respectively[1, 10], as detailed in Methods of main text. Random effects for $\log_{10}\beta'$ and $\log_{10}T(0)$ were less than 0.1, and were thus removed, rendering them to be same across macaques.

| Parameter | Description | Value | Reference |
|---|---|---|---|
| $\hat{\beta}$ | Infection of target cells by virions | $10^{-8}$ mL cells$^{-1}$ d$^{-1}$ | [11] |
| $\delta$ | Death of virus-producing cells | 1.44 d$^{-1}$ | [12] |
| $\rho$ | Transition from eclipse phase to actively producing virions | 0.36 d$^{-1}$ | Estimated; Fig. S1a |
| $\hat{p}$ | Virus production | 1440 d$^{-1}$ | [12] |
| $c$ | Free virion clearance | 0.35 d$^{-1}$ | [12] |
| $f$ | Fraction of infections resulting in non-productive infections | 0.5 | [13] |
| $\hat{T}_0$ | Initial number of CD4 T-cells | $10^6$ cells mL$^{-1}$ | [6] |
| $\hat{V}_0$ | Viral inoculum size | $10^{2.86}$ mL$^{-1}$ | [6] |
| $\hat{C}_0$ | Concentration of CD8 T-cells extracted for the *ex vivo* assay | $10^6$ cells mL$^{-1}$ | [6] |
| $C_0$ | Concentration of total CD8 T-cells in infected hosts | $10^6$ cells mL$^{-1}$ | [14] |
| $\mathcal{R}_0$ | Basic reproductive ratio of virus in cultures | 14.29 | Estimated |
| $\epsilon$ | Fraction of target cells infected by peak viral load in CD4 T-cell culture | 0.93 | Estimated |
| $\tau_{max}$ | Time point in the CD4 T-cell culture when the viral load peaks | 5.64 d | Estimated |

**Table S7: Parameters of the *ex vivo* model.** The table lists the values used, and the references thereof. CD8 T-cell count in untreated SIV-infected cynomolgus macaques was close to $10^6$ cells mL$^{-1}$ (ref [14]), similar to the levels in HIV-infected humans[15, 16]. So, we fixed $C_0$ to $10^6$.

| Parameter (Units) | Fixed effect | Random effect |
|---|---|---|
| $\lambda$ (cells mL$^{-1}$ d$^{-1}$) | 0.03 (0.06) | 2.00 (1.87) |
| $\log_{10}\beta'$ (log mL cells$^{-1}$ d$^{-1}$) | -1.73 (0.06) | 0.04 (0.03) |
| $f_D$ | 0.95 | - |
| $\log_{10}\lambda_E{}^*$ (log d$^{-2}$) | -0.29 (0.02) | 0.01 (0.01) |
| $d_I$ (d$^{-1}$) | 0.1 | - |
| $d_D$ (d$^{-1}$) | 0.01 ($3.04\times10^{-3}$) | 1.00 (0.33) |
| $\gamma$ (cells$^{-1}$) | $7.42\times10^{4}$ ($2.77\times10^{4}$) | 1.35 (0.28) |
| $\alpha_E$ (d$^{-1}$) | 0.62 (0.02) | 0.09 (0.02) |
| $\theta_E$ (cells mL$^{-1}$) | 0.10 | - |
| $d_E$ (d$^{-1}$) | 1.00 | - |
| $\log_{10}T(0)$ (log cells mL$^{-1}$) | 3.38 (0.06) | 0.01 (0.01) |

**Table S8: Population parameter estimates for model which does not incorporate suppressive capacity measurements.** The fixed and random effects of each parameter are provided along with respective standard errors in parentheses. In addition to the parameters fixed in the best-fit model, $f_D$ is fixed to 0.95 (ref [9]).

| **Variable** | **Description** | **Initial value** |
|---|---|---|
| $T(0)$ | Initial concentration of target CD4 T-cells | $\lambda/d_T$ |
| $I(0)$ | Initial concentration of productively infected cells | 0 |
| $D(0)$ | Initial concentration of non-productively infected cells | 0 |
| $V(0)$ | Viral inoculum size | $10^{-2.76}$ copies $mL^{-1}$ for macaques BC094, BC179, BD536, and BO186<br>$10^{-1.76}$ copies $mL^{-1}$ for the other macaques |
| $E(0)$ | Initial concentration of SIV-specific effector CD8 T-cells | 0 |
| $k(0)$ | Killing rate constant of productively infected cells by virus-specific CD8 T-cells | 0 |
| $Q(0)$ | Initial exhaustion level | 0 |

**Table S9: Initial conditions used for *in vivo* model fitting.** Note that the exhaustion compartment, *Q*, is present only in models #3 and #4. Viral inoculum sizes have been estimated using the volumes of distribution[17].
